# Green notes: The rhythms of cyanobacteria exoelectrogenesis as de-composed by the Hilbert-Huang transform

**DOI:** 10.1101/2021.10.22.465414

**Authors:** Tonny Okedi, Kamran Yunus, Adrian Fisher

**Affiliations:** Department of Chemical Engineering and Biotechnology, University of Cambridge, Phillipa Fawcett Drive, Cambridge CB3 0AS, UK; Cambridge Center for Advanced Research and Education in Singapore (CARES), 1 Create Way, #05-05 CREATE Tower, Singapore 138602, Singapore

## Abstract

Electrons from cyanobacteria photosynthetic and respiratory systems are implicated in current generated in biophotovoltaic (BPV) devices. However, the pathway that electrons follow to electrodes remains largely unknown, limiting progress of applied research. Here we use Hilbert-Huang transforms to decompose *Synechococcus elongatus* sp. PCC7942 BPV current density profiles into physically meaningful oscillatory components, and compute their instantaneous frequencies. We develop hypotheses for the genesis of the oscillations *via* repeat experiments with iron-depleted and 20% CO_2_ enriched biofilms. The oscillations exhibit rhythms that are consistent with the state of the art cyanobacteria circadian model, and putative exoelectrogenic pathways. In particular, we observe oscillations consistent with: rhythmic D1:1 (photosystem II core) expression; circadian-controlled glycogen accumulation; circadian phase shifts under modified intracellular %ATP; and circadian period shortening in the absence of the iron-sulphur protein LdpA. We suggest that the extracted oscillations may be used to reverse-identify proteins and/or metabolites responsible for cyanobacteria exoelectrogenesis.

## 2 Introduction

There is a critical need for CO_2_ capture and abatement technologies in order to meet global climate change goals for less than 2°C of warming. Biophotovoltaic systems (BPVs) which employ microorganisms that perform oxygenic photosynthesis have emerged as one such potential technology. A BPV is an electrochemical cell in which at least one electrode is catalysed by photosynthetic microorganisms such as algae and cyanobacteria, that absorb CO_2_ from the atmosphere, reduce it into carbon products during photosynthesis, and store it as biomass. Electrons generated from the photosynthetic process have been implicated in the enhanced current generated in these devices under illumination (photo-current), and in the basal current in the dark (dark current) from oxidation of the stored biomass *e*.*g*., *via* respiration or the oxidative pentose phosphate pathway ^1–4^.

Key to delivering BPVs is understanding why the microorganisms donate electrons to their surroundings (exoelectrogenesis) and the complex electron flows from the photosynthetic and respiratory electron transport chains, to the non-living electrodes, Fig. 1c. This remains a major hurdle in advancing the fundamental understanding needed to develop more efficient light conversion, and to enable applied research to realise the technology at a commercial scale.

**Figure 1.**
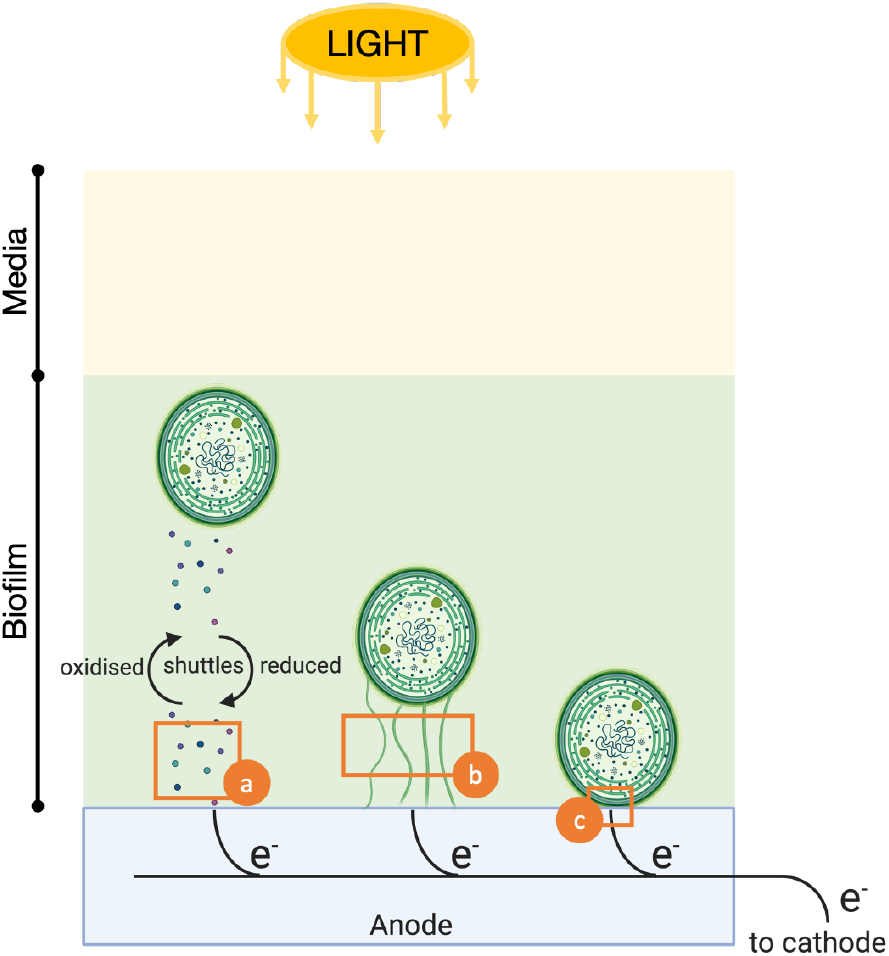
Putative terminal electron transport step in cyanobacteria. (**a**) indirect electron transport *via* endogenous electron mediators excreted by the cells; (**b**) direct electron transfer *via* extracellular appendages that traverse the cell cytoplasmic membrane and outer cell wall in contact with the electrode; and (**c**) direct electron transfer *via* redox active proteins embedded in the cell outer membrane. Schematic is not to scale.

To date, electron flows have been primarily investigated *via* spectroscopy and electrochemical techniques such as chronoamperometry and cyclic voltammetry studies on mutant, stressed or inhibited cells ^2–8^. Computational models and tools are being developed that can be used to help interpret experimental data and run rapid simulations towards understanding the electron flows ^9–11^. However, due to knowledge gaps, the non-linear and nonstationary nature of bioelectrochemical processes, and difficulties in the in-situ measurement of key variables, it has proved difficult to develop robust first-principle computational models. To overcome these challenges, the authors recently applied deep learning, in particular Long Short-Term Memory (LSTM) networks, to predict the one-step-ahead seasonal current density in BPVs using a three step process: (1) decomposing the observed current density into a trend, seasonal and irregular component *via* Seasonal and Trend Decomposition using Locally Estimated Scatterplot Smoothing or LOESS (STL); (2) training the LSTM network with the seasonal component; and (3) predicting the one-step-ahead seasonal current density using the trained network ^12^.

While satisfactory results were achieved, the LSTM model was limited to predicting the repeatable seasonal current density. In order to forecast the observed current density, a separate model for the erratic and difficult-to-predict irregular component is required. The LSTM forecasts were shown to be unsatisfactory without the removal of the irregular component^12^. Furthermore, the physical meanings of the trend, seasonal and, particularly, irregular components obtained *via* STL are unclear. It is therefore imperative to find an alternative decomposition approach that results in more regular, physically meaningful sub-components in order to develop more robust models *via* LSTMs or other time series modelling techniques. To achieve this, the authors previously proposed Empirical Mode Decomposition (EMD), a time series decomposition method applied when analysing signals using the Hilbert-Huang transform (HHT) ^12^.

The Hilbert-Huang transform is a two-step time series analysis technique for non-linear and/or nonstationary signals such as exoelectrogenic currents^13^. The first step is a pre-processing EMD stage. The goal of EMD is to expand a nonstationary signal made up of multiple, superimposed, dynamic oscillations, into a finite set of sub-components or intrinsic mode functions (IMFs). Each IMF (or simply mode) includes oscillations of a characteristic, physically meaningful, timescale (meaning the time between oscillatory peaks) ^13^. This allows the Hilbert transform to be applied to each IMF to obtain meaningful instantaneous frequencies along its time course. Thus, HHT is an amplitude-frequency-time technique that allows the identification of events on both the time and frequency axes.

It was hypothesised that by applying the HHT to current density profiles from unmediated BPVs, distinct oscillatory patterns can be isolated, from which characteristic timescales and frequencies intrinsic to natural exoelectrogenesis may be identified. In this work, the decomposition step is implemented with a modified EMD algorithm called Improved Complete Ensemble EMD with Adaptive Noise (ICEEMDAN), developed to result in more physically meaningful IMFs (see Methods for more details) ^14^. To help decode the physical meaning of the extracted IMFs, BPVs were operated in three different conditions known to affect exoelectrogenesis in distinct ways: (1) standard laboratory growth conditions (control); (2) iron depleted media; and (3) iron depleted media in the presence of 20% CO_2_. BPVs operated with iron-depleted *Synechococcus elongatus* sp. PCC7942 (*S. elongatus* henceforth) biofilms exhibit significantly larger currents but a markedly reduced light response ^7,15^. Peculiarly, BPVs operated with iron-depleted cultures in the presence of 20% v/v CO_2_ and with ferricyanide as an exogenous mediator, exhibit higher current output in the dark than in the light^15^. The enhanced currents under iron starvation was suggested as evidence of the over-expression of a redox-active protein involved in iron acquisition at the outer membrane level ^7^. The enhanced dark current in the presence of 20% CO_2_ was suggested to be evidence of preferential utilisation of stored energy for exoelectrogenesis.

Throughout the paper, Fe (+)|Air refers to the control condition (black lines in graphs), Fe (-)|Air refers to iron depleted cultures/biofilms in atmospheric air (green lines in graphs), and Fe (-)|20% CO_2_ refers to iron depleted cultures/biofilms in the presence of 20% CO_2_ (blue lines in graphs). The additional suffix or subscript |3h refers to operation under a 3h:3h light-dark period, while |12h refers to a operation under a 12h:12h (diel) light-dark period. Finally, borrowing from terminology used in chronobiology, *dusk* refers to light-to-dark transitions, *dawn* to dark-to-light transitions, *day* to the illuminated interval of a period, and *night* the dark interval.

## 3 Results

### 3.1 Current density profiles

Figure 2 shows the measured current density profiles for the three conditions investigated following the operating procedure in Tab. 2 (Methods). Current densities from abiotic BPVs inoculated with the corresponding fresh medium for each condition are also shown to give an indication of the background current. It should be noted that the magnitudes of the backgrounds are exaggerated because in the biotic devices, the cells assimilate redox active species in the medium over time, reducing their concentration. This is particularly true for the Fe (+)|Air condition, where redox active ferric ions are assimilated by the cells. We previously estimated that the concentration of ferric ions in fresh BG11 medium reduces by half within four days of culturing and reduces to almost zero within 11 days under similar growth conditions ^16^. Similar rates of ferric ion depletion rates in *S. elongatus* cultures have been independently reported from atomic absorption spectroscopy concentration measurements ^17^.

**Figure 2.**
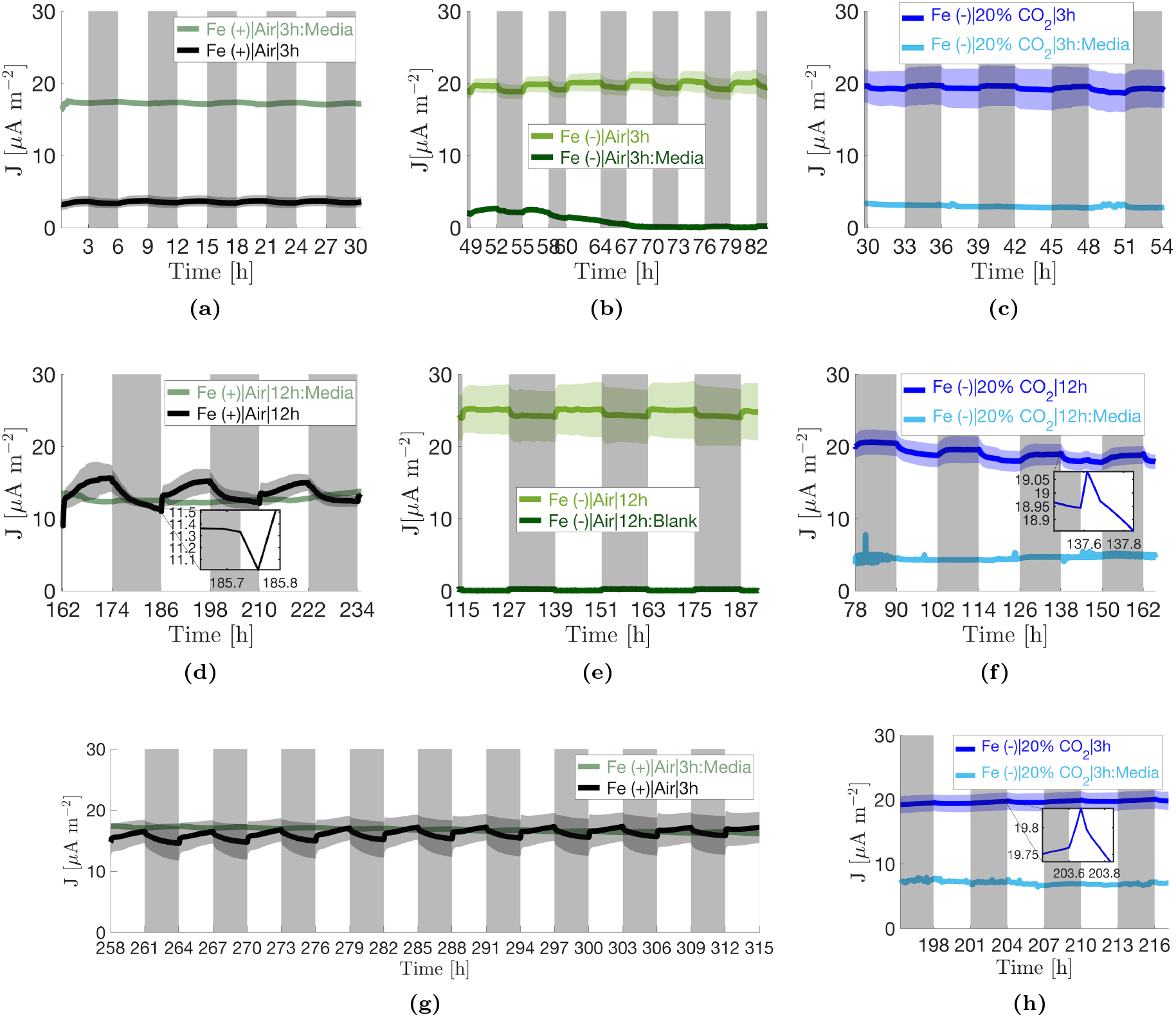
Current density profiles. Each profile shows the mean of three independent replicates ± 1 standard error of the mean (shaded areas). Current profiles of abiotic BPVs inoculated with the corresponding fresh media are also shown. BPVs were operated with the procedure shown in Tab. 2. Profiles **(a)–(c)** and **(g)–(h)** show operation under 3h:3h light-dark periods while **(d)**–**(f)** show operation under 12h:12h light-dark periods. Profiles (g)–(h) were recorded after replenishment of respective media into the BPVs. The time gaps between the corresponding 3h:3h profiles and the 12h:12h profiles includes *≈* 10h during which polarisation curves where measured (Fig. S10), and the remaining time the duration of voltage recovery and stabilisation following reconnection of the 33 *M*Ω external resistors. This duration varied by operating condition. In (**c**), a light timer error resulted in a two-rather than three-hour dark interval at 58h, and consequently a four-rather than three-hour light interval ending at 64h.

#### Non-stressed cells exhibit a two-sloped exoelectrogenic signal after a ca. 40h time lag

The Fe (+)|Air BPVs did not produce significant current during the first 30h of operation under load, Fig. 2a. After measurement of a polarisation curve in the dark (Fig. S10a) and reconnection of the 33 *M*Ω external resistor, the current densities increased erratically from around 40h and began to stabilise at around 140h (data not shown). Repeatable oscillations were observed in all three replicates after approximately 162h, Fig. 2d. The time scales of the different current generation phases (negligible, erratic, stable) mirror the evolution of Fe (+)|Air culture growth from lag to stationary phase (Fig. S2). It is therefore hypothesised that: (i) the negligible current density in the first 30h is due to cells in the lag phase prioritising accumulation of resources, including reducing power, for cell proliferation; (ii) the erratic increase in current density is due to biofilms in exponential phase, where there is rapid growth in electrochemically active biomass in the biofilm; and (iii) the stable currents are achieved when the biofilm reaches stationary phase where concentration of active biomass in the biofilm remains constant, and gradual changes in cellular morphology contribute to changes in the mean current level by influencing electron export rates ^16^. Patterns previously reported when operating under a diel (24h) light-dark period were observed, namely a dip and recovery in current density at dawn (inset, Fig. 2d), and a period-to-period decay in the magnitude of the photocurrent^12^. Following the dip, photoresponse was two-sloped, with an initial steep spike in current, followed by a more gradual increase over the remaining duration of the illuminated interval. In addition, the slope of the initial spike decayed substantially with each period, dropping from around 20 *μA m*^*−*2^ *h*^*−*1^ at the 162h dawn to around 6.5 *μA m*^*−*2^ *h*^*−*1^ at the 234h dawn, Fig. S11d.

To test whether the decaying photoresponse in the Fe (+)|Air BPVs was due to the depletion of the Fe (+)|Air medium, BPVs were replenished with fresh BG11 media, reconnected to 33 *M*Ω resistors and operated with a 3h:3h light-dark period, Fig. 2g. The mean current density maintained its level, and even increased slowly over time. The slope of the initial positive photoresponse also showed significant recovery reaching approximately 13 *μA m*^*−*2^ *h*^*−*1^ at the 295h dawn, Fig. S11g. However, the second, gradual photoresponse was not recovered, and overall photoresponse remained low. The dip in current at dawn was absent under the 3h:3h light-dark regime confirming previously reported results ^12^.

#### Iron stress expedites activation of exoelectrogenesis, amplifies mean current output, but reduces light gradients

In line with previous results, Fe (-)|Air BPVs had higher current densities versus the control, but a markedly reduced photoresponse, Fig. 2b and e. Current densities increased erratically for the first 40h-45h upon connection to a 33 *M*Ω external resistor (data not shown). All 3 replicates produced stable and repeatable current oscillations from 49h onwards. This was substantially quicker than in the control. Similar to the Fe (+)|Air devices, the evolution from erratic to stable current output mirrors the evolution of Fe (–)|Air culture growth *i*.*e*., a substantially shorter lag phase (< 6 h) and transition to decline within 45 h of inoculation (Fig. S2 and Tab. S2). This further corroborates the hypothesis that stable currents are achieved when biofilms reach the stationary phase. At 82h, the light-dark period was switched from 3h:3h to 12h:12h. It took approximately 33 hours to reestablish stable and repeatable oscillations which were achieved in all three replicates from 115h onwards, Fig. 2e. This was substantially quicker than in the control. The mean current density also increased slightly during this time. Unlike in the Fe (+)|Air|12h BPVs, there was no dip in current at dawn and photoresponse was mono-sloped. The photocurrent therefore plateaued after the initial rise. As in the control, the slope of the photoresponse decayed from period to period, albeit at a much slower rate. The slope was also less steep, reducing from approximately 4 *μA m*^*−*2^ *h*^*−*1^ at the 49h dawn, to 2 *μA m*^*−*2^ *h*^*−*1^ at the 187h dawn, Fig. S11b and e.

#### The presence of 20% CO_2_ further accelerates the start of exoelectrogenesis and amplifies the dark current

Fe (-)|20% CO_2_ BPV current densities were similar in magnitude to those of the Fe (-)|Air condition, Fig. 2c and f. Distinctively, the current was marginally higher in the dark than in the light, with a conspicuous lack of the enhanced photocurrent typical in BPVs. The results confirm observations previously reported in the research group, here also observed in unmediated devices ^15^. Counterintuitively, there was a positive dark response at dusk, with a diminishing slope of less than 1.8 *μA m*^*−*2^ *h*^*−*1^, Fig. S11c and f. With every subsequent dusk, the dark response became less prominent. Oppositely, an ephemeral positive photoresponse at dawn appeared over time, first seen at 114h and slowly increased in prominence with each new dawn, inset of Fig. 2f. The Fe (-)|20% CO_2_ BPVs achieved stable and repeatable oscillations within the first 30h of connection to a 33 *M*Ω external resistor, Fig. 2e. Following the polarisation curve at 54h, the BPVs reestablished stable and repeatable oscillations within 12-14h of reconnecting the 33 *M*Ω external resistor, much faster than the other two conditions, Fig. 2f. This is consistent with the rapid evolution of Fe (–)|20% CO2 cultures to stationary phase (Fig. S2 and Tab. S2).

To check the persistence of the ephemeral positive photoresponse that first appeared at the 114h dawn, the Fe (-)|20% CO_2_ BPVs were operated for a further 21 hours under a 3h:3h light-dark period after the light polarisation at 162h, Fig. 2h. The persistence of the photoresponse was confirmed as clearly seen in the derivative of the current density profile, Fig. S11h. Further, it was observed that in the long term, the current profile became flatter and flatter as the positive dark response continued to diminish.

### 3.2 Hilbert-Huang transforms

Each of the profiles in Fig. 2, were decomposed using the ICEEMDAN algorithm (see algorithm 2 under Methods), resulting in the intrinsic mode functions displayed in Fig. 3 and Fig. 6a-b. The Hilbert transform was then performed on all the extracted IMFs to compute the instantaneous frequencies and energies. The data from the computed analytic signals can be displayed as frequency profiles contoured with instantaneous energies such as displayed in Fig. 4 for the Fe (-)|Air|3h condition. When plotted on a single axis, the contoured instantaneous frequency profiles form the Hilbert spectrum. The Hilbert spectra for the different operating conditions are shown in Fig. 5 and Fig. 6c-d. In addition, the IMFs and Hilbert spectra of the corresponding abiotic, media-only devices are shown in Figs. S12–13 to identify the background. For some modes, both the peak-to-peak duration and median periodicity (computed from the inverse of the median instantaneous frequency shown in Fig. 5) are quoted. Reason for discrepancies in the two numbers is addressed in the Discussion section of the paper.

**Figure 3.**
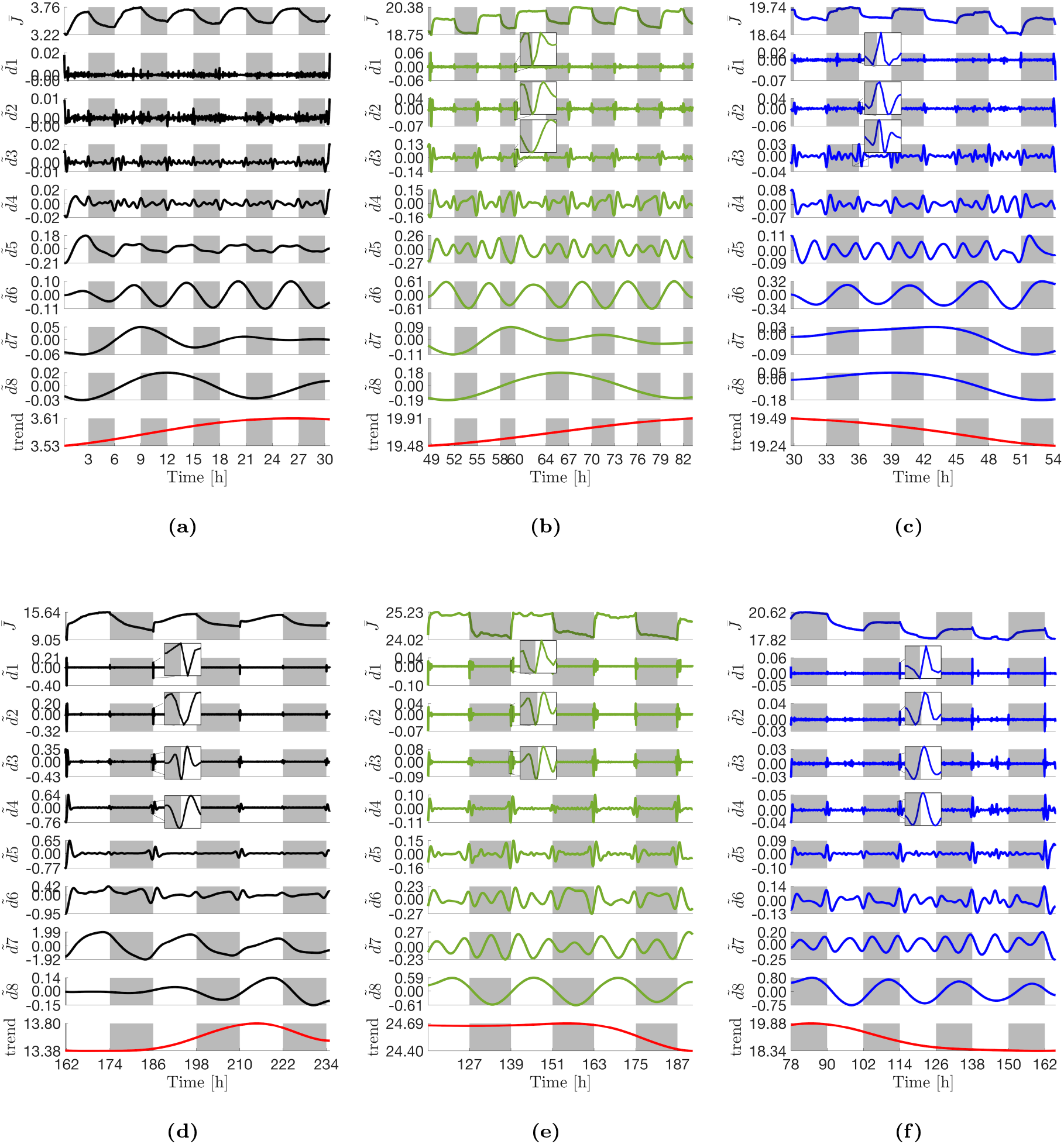
Intrinsic mode functions (IMFs) extracted *via* the ICEEMDAN algorithm. **(a)** Fe (+)|Air|3h. **(b)** Fe (-)|Air|3h. **(c)** Fe (-)|20% CO_2_|3h. **(d)** Fe (+)|Air|12h. **(e)** Fe (-)|Air|12h. **(f)** Fe (-)|20% CO2|12h. For each decomposition, the top panel shows the mean current density profile, 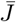, as reported in Fig. 2, the central panels 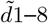 show the extracted IMFs, and the bottom *trend* panel shows the final residue from the decomposition process *r*8 (red line). Insets in (**b**)-(**f**) show zooms of the highest frequency oscillations at dawn for improved clarity. It should be noted that the widths and heights of the insets reflect varying durations (increasing as you move downward from 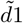) and amplitudes. Light changes are superimposed as the backgrounds of the panels (light on white and light off grey).

**Figure 4.**
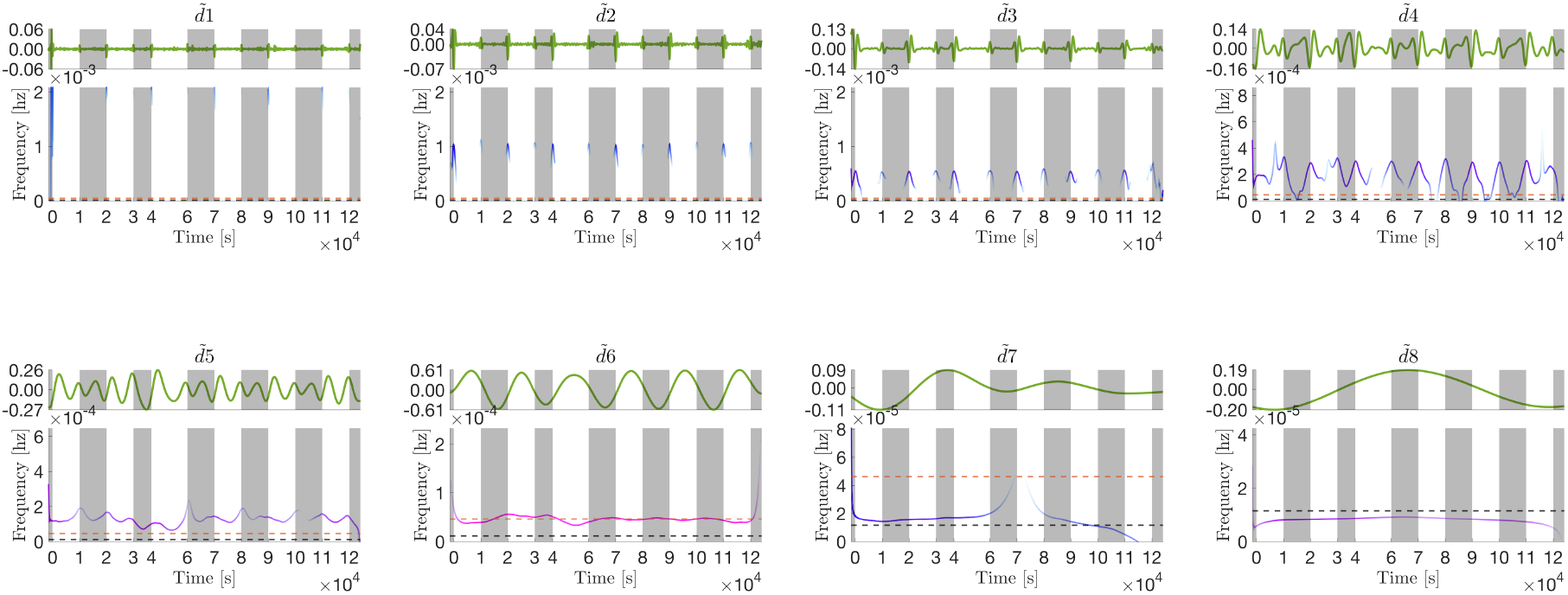
IMFs in Fig. 3b extracted from the Fe (-)|Air|3h current density profile (top panes) and their corresponding instantaneous frequency profiles (bottom panes). The instantaneous frequency profiles are contoured with the instantaneous energy of the oscillations (see Fig. 5 for the colour bar). The orange dashed line marks the frequency of the light-dark period, while the black dashed line marks the diel (24h) frequency. When plotted on one axis, the contoured frequency profiles above form the Hilbert spectrum displayed in Fig. 5c. Note the x-axes origins have been zeroed and time is shown in seconds (standard representation).

**Figure 5.**
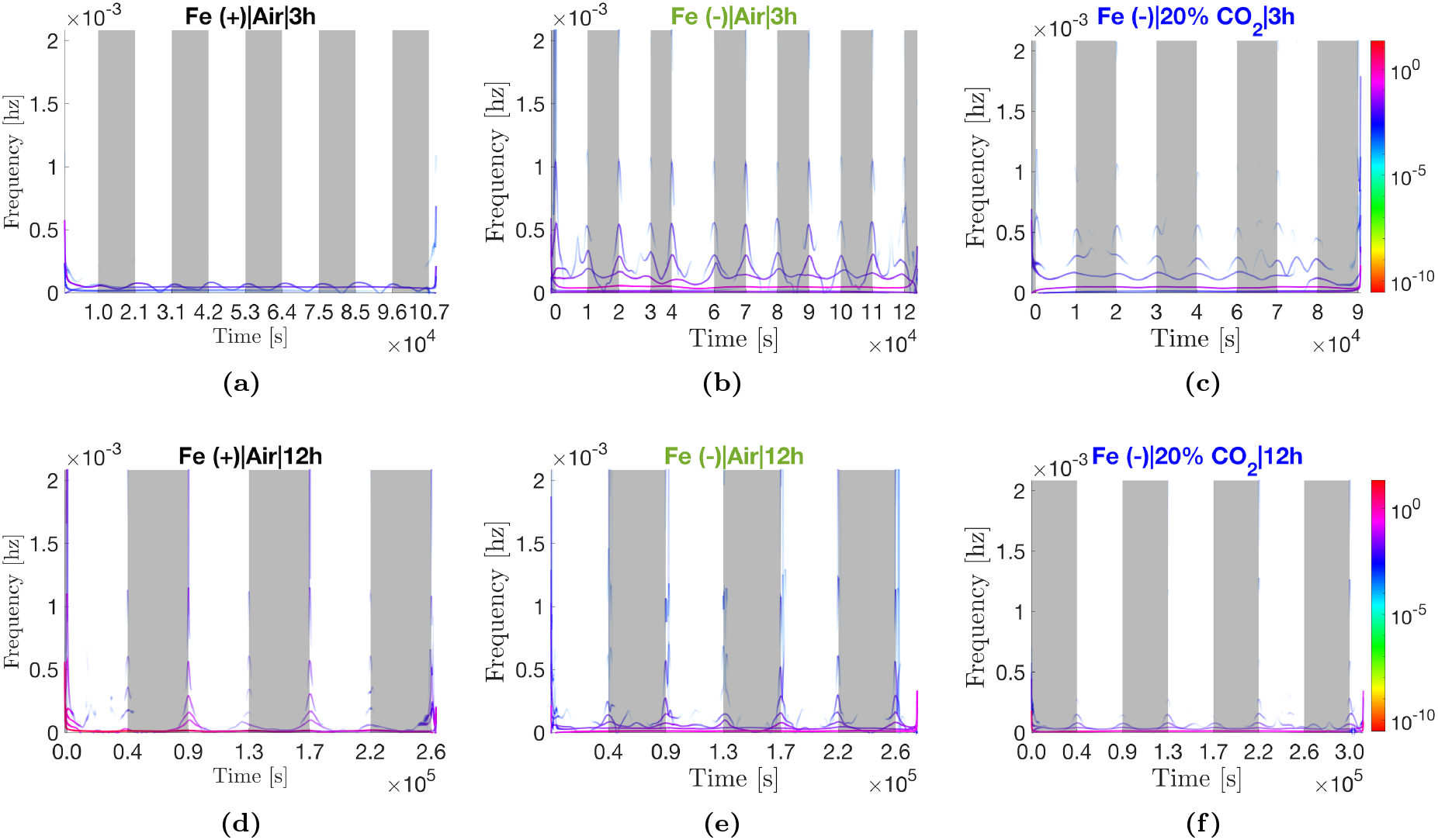
Hilbert spectra of the different current density profiles showing the energy-frequency-time distributions. Each spectrum is a single-axis plot of the full set of contoured instantaneous frequency profiles computed from the IMFs of the corresponding operating condition (Fig. 3). For example, **(c)** shows the single-axis plot of the instantaneous frequency profiles shown in Fig. 4. The y-axis is limited to the maximum frequency than can be resolved with a two-minute sampling interval (see Eq. 7). Note the x-axes origins have been zeroed and time is shown in seconds (standard representation of the Hilbert spectrum).

**Figure 6.**
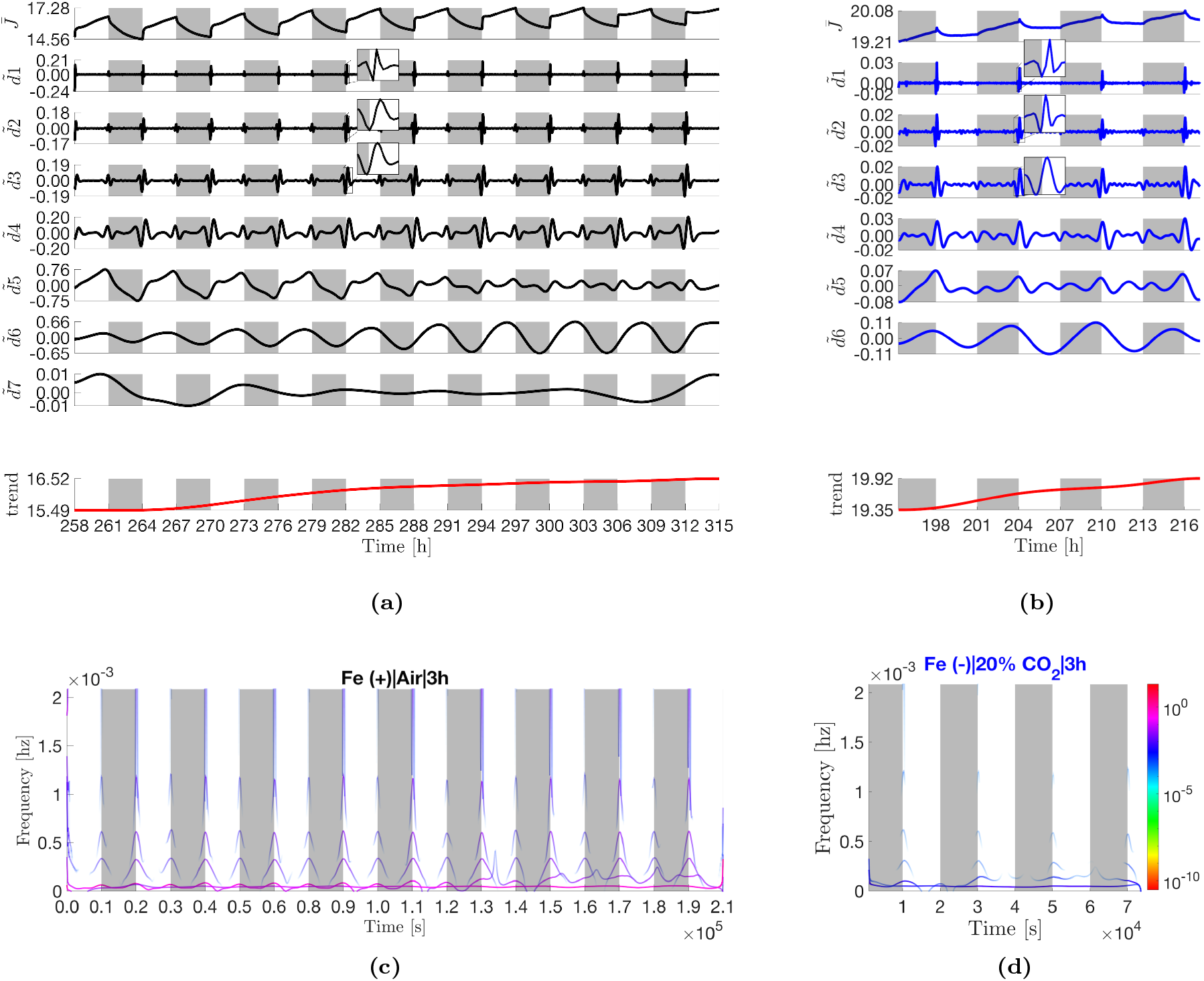
IMFs and Hilbert spectra of Fe (+)|Air|3h and Fe (-)|20% CO_2_|3h after replenishment of respective media.

### Eight modes are extracted from the control current density profile

Eight IMFs were extracted from the Fe (+)|Air|3h current density profile, Fig. 3a. Signals in the first four IMFs, 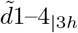, are indistinguishable from measurement noise - no oscillations with significant energies (amplitudes) or frequencies were observed. It will be shown later that conspicuous oscillations with amplitudes significantly higher than noise develop at dawn in these modes. IMF 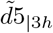 is a persistent frequency and amplitude modulated (FM-AM) mode. The mode exhibits a repeating pattern of an intermediate peak in the early day (*≈*1h after dawn), a high peak just before dusk and a low peak in the night. The peak-to-peak timespans are 1.51±0.04h (1^st^ to 2^nd^ peak) and 2.17±0.14h (2^nd^ to 3^rd^ peak). Maximum positive frequency deviations occur at dusk and at dawn, while maximum negative frequency deviations occur between the two light transitions, Fig. 5a. IMF 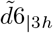 oscillates with a peak-to-peak time span of 5.9±0.04h and a median period of 6.0h, and peaks around 75% into the day. Interestingly, despite the imposed 6h (3h:3h) light-dark periodicity, IMFs 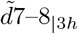 oscillate with constant periods of approximately 13.1h and 20.6h, respectively.

#### Iron stress triggers oscillations that only developed later on in the control

The Fe (-)|Air|3h current density profile also expanded to eight IMFs, all exhibiting larger amplitudes than their respective control (Fe (+)|Air|3h) analogs, Figs. 3b and 5b. Notably, prominent oscillations with amplitudes appreciable larger than the measurement noise are visible at dawn in IMFs 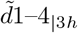, Fig. 4. These oscillations will be shown to only develop later in the control. All 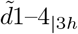 oscillations have their minima at dawn, and their energies and frequencies fall to zero following a single cycle *i*.*e*., they are impersistent. While the FM-AM mode 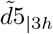 also exhibits the same repeating three-peak pattern as its control analog, the amplitude ratios and timings are altered. Under iron starvation, the highest peak is the early day peak (now just *≈*0.6-0.7h after dawn) and the dusk-timed and nighttime peaks are of roughly equal amplitude that is lower than that of the early day peak. The 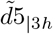 peak-to-peak timespans are 2.19±0.02h (1^st^ to 2^nd^ peak) and 1.89±0.02h (2^nd^ to 3^rd^ peak). Mode 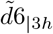 had 6.0±0.1h peak-to-peak timespan and 6.0h median period but is phase shifted by approximately *π/*4 rads towards dawn, resulting in oscillations peaks earlier in the day (approximately halfway through). Finally, 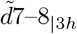 have periods of 16.1h and 33.2h, respectively, both significantly longer relative to the control.

#### All modes exhibit an apparent *π* rad phase shift and reduced amplitudes in the presence of 20% CO_2_

The Fe (-)|20% CO_2_|3h current density profile expanded to eight IMFs, Fig. 3c. The oscillations of 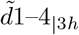 have approximately the same frequencies as their respective control and Fe (-)|Air|3h analogs, albeit generally lower energies than the latter, Fig. 5c. As expected from the current density profile (Fig. 2c), all IMFs have a *π* rad phase shift relative to the control, most clearly exemplified in 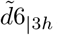, Fig. 3c. The FM-AM mode 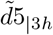 has similar peak-to-peak timespans as its Fe (-)|Air|3h analog, 2.14±0.03h (1^st^ to 2^nd^ peak) and 1.93±0.13h (2^nd^ to 3^rd^ peak). Mode 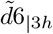 has a peak-to-peak timespan of 6.0±0.4h and a median period of 6.0h. Modes 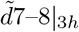 oscillate with longer periods than their control analogs at 25h and 27h, respectively. However, there is uncertainty due to an abnormally prolonged decline in current density between 48–51h of operation that caused mode mixing (see methods for explanation) across 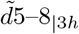 (visible from 48–54h).

### The superposition of the four highest frequency modes is responsible for the initial steep photoresponse

There are some notable differences in the control (Fe (+)|Air) IMFs extracted from the 12h:12h current density profiles compared to the 3h:3h regime. First, there is a large increase in the amplitudes of all oscillations in the Fe (+)|Air|12h IMF set. As a result, conspicuous single-cycle oscillations are visible at dawn in 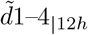, Figs. 3d and 5d. These oscillations have the same frequencies as 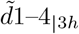 extracted from the Fe (-)|Air|3h current density profiles, but with significantly larger amplitudes. In the non-stressed condition, the oscillations are only triggered after >59h vs. within 30h in the iron stressed biofilms. IMF 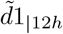 oscillations peak at dawn, while in 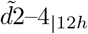, the peaks are increasingly shifted to the left of dawn, with 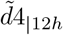 having minima coincident with the light transition, insets Fig. 3d. The superposition of 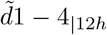 is thus revealed as responsible for the dip in current density observed immediately after illumination and the ensuing steep recovery segment of the two-sloped photoresponse, inset Fig 2d. The amplitudes of 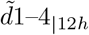 oscillations decrease with each successive dawn, accounting for both the shallower dip and the diminishing gradient of the initial photoresponse from period-to-period.

### Two distinct modes revealed under 12h:12h light-dark periodicity

Secondly, two modes, 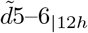, with distinct frequencies absent from the Fe (+)|Air|3h IMF set are extracted under the 12h:12h regime, Fig. 3d. IMF 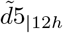 contains single-cycle oscillations with minima at dawn while IMF 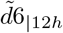 is a persistent FM-AM mode. Both 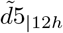 and 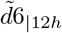 contribute to the first of the two-sloped photoresponse. IMFs 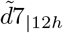 and 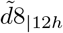 are analogs of the FM-AM 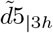 and the constant frequency 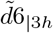 in the 3h:3h regime, respectively, Fig. 3d. Mode 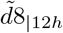 oscillates with peak-to-peak duration of 24.0±1.1h and a median periodicity of 25.8h. In comparing 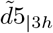 (Fig. 3a) and 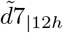 (Fig. 3d), it can be seen that a large increase in the amplitude of the dusk-timed peak, with a concomitant decrease in the amplitude of the nighttime peak occurred during the 40-162h interval. In contrast, the change in the amplitude of the early day peak is marginal. The relative changes in amplitude drastically increase the prominence of the second peak such that the early day and nighttime peaks appear instead as inflection points. The changes reveal IMF 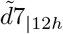 as responsible for the second gradient of the two-sloped photoresponse seen in the current density profile. Over time (162-234h), there is a redistribution of the energies - the amplitude of the dusk-timed peak decreases with a concomitant increase in the amplitude of the nighttime peak, accounting for the dwindling photocurrent observed in Fig 2d. It will be shown in the decomposition of the Fe (-)|Air|12h that peak-to-peak durations of 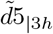 and 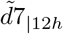 are linearly scaled to the duration of the imposed light-dark period. Thus, despite the hidden peaks in 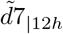, the peak-to-peak timespans are estimated as *≈*6.04±0.16h (1^st^ to 2^nd^ peak) and *≈*8.68±0.56h (2^nd^ to 3^rd^ peak) by linearly scaling the analogous 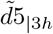 timespans. Interestingly, modes 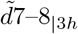 from the Fe (+)|Air|3h decomposition, seemingly disappear when BPVs are operated under a 12h:12h light-dark pattern.

#### Iron stress shortens the periods of persistent modes by 1.2h under 12h:12h light-dark periodicity Iron

stress (Fe (-)|Air|12h) caused a phase shift such that the peaks advance towards dawn which is most visible in 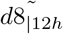, Fig. 3e. In 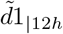, the phase shift results in the minima, rather than the maxima, of oscillations occurring at dawn, inset Fig. 3e. The absence of a current dip after illumination in the Fe (-)|Air|12h BPVs is attributable to this change. Under a 12h:12h light-dark period, the peak-to-peak timespans of the FM-AM 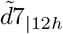 are linearly scaled to 8.61±0.36h (1^st^ to 2^nd^ peak) and 7.43±0.15 (2^nd^ to 3^rd^ peak). Notably, the duration between 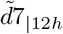 dusk-timed and nighttime peaks, as well as the peak-to-peak duration and median periodicity of 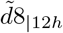 (23.1±1.1h and 24.6h, respectively) are shortened by 1–1.2h relative to the control. The duration between 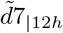 early day and dusk-timed peaks increases by *≈* 2.5*h* due to an advance of the early day peak towards dawn under iron stress.

#### Amplitude changes in the impersistent modes are correlated with ferric ion concentration in the media

In comparing Figs. 3d and 3e, it can be seen that, the amplitudes of 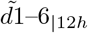 oscillations in the control diminish across successive periods until, at 234h, they reach similar magnitudes as their Fe (-)|Air|12h analogs. Given that the only difference in the two conditions was the initial ferric ion concentration in the respective growth media, the results hint that the decaying amplitudes are a function of gradual ferric ion depletion in the Fe (+)|Air medium. To corroborate this conclusion, amplitudes of 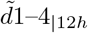 analogs, 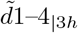, were observed to be ferric-sensitive. That is, when the depleted media in the Fe (+)|Air BPVs were replenished with fresh BG11 at *≈*246h and the BPVs operated under a 3h:3h light-dark period (Fig. 6a), the modes exhibited period-to-period recoveries in amplitude until the 294h dawn, before decaying again, Fig. 6a. Modes 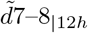 are exceptions to this. Amplitude in 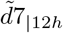 analog 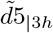 did not necessarily recover after ferric ion replenishment. Rather the amplitude decay (particularly of the dusk-timed peak) was arrested until ca. 282h when appreciable decay restarted. By 288h, the three distinct peaks are clearly visible as the prominence of the dusk-timed peak continues to diminish and by 306h, the early day peak becomes the highest peak as in the Fe (-)|Air condition. The 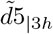 peak-to-peak timespans also evolve to match those of the Fe (-)|Air|3h condition, 2.15±0.07h (1^st^ to 2^nd^ peak) and 1.78±0.09h (2^nd^ to 3^rd^ peak). Oppositely, the amplitude of 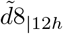 increases with time to a similar magnitude as its Fe (-)|Air analog by 288h, Fig. 3d and 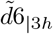 in Fig. 6a.

#### Positive light response in the presence of CO_2_ is due the four highest frequency modes

As in the 3h:3h regimes, the presence of 20% CO_2_ caused a *π* rad phase shift in the Fe (-)|20% CO_2_|12h IMFs relative to the 12h:12h control, Fig. 3f. The oscillations also exhibited the diminished 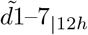 amplitudes observed under iron depletion. Mode 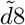 had a peak-to-peak timespan of 24.3±0.7h and a median period of 24.4h. With the exception of 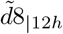, amplitudes were even lower than their Fe (-)|Air|12h analogs. Relative to 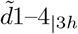 (Fe (-)|20% CO_2_|3h, Fig. 3c), small phase differences are visible in 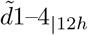 that push the peaks of the oscillations just to the right of dawn, insets Fig. 3f. A period-to-period increases in amplitudes of the modes is also visible. With these changes, it becomes evident that the superposition of 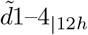 form the short-lived positive light response observed beginning from the 114h dawn, inset Fig. 2f. Under a 12h:12h light-dark period, the peak-to-peak timespans of the FM-AM 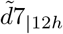 are scaled to 8.17±0.14h (1^st^ and 2^nd^ peaks) and 8.02±0.18 (2^nd^ and 3^rd^ peaks).

#### Iron stress increases the basal exoelectrogenic activity

The effect of iron starvation is most pronounced in the residuals, *i*.*e*. the trends, Fig. 3. In the early stages of operation (<80h), there was an up to 5.7 fold increase in the magnitude of the residual under iron depleted conditions, which declines to approximately 1.8 fold in the later stages (>162h) of operation as ferric ions are also depleted in the control BPVs. The result that iron starvation increases the basal exoelectrogenic activity in *S. elongatus* is consistent with previous results reported by the research group ^7^.

## 4 Discussion

In the following discussion, initial hypotheses on the physical meaning of the modes are developed for future investigations. The first clue available in the results in the 25.8h median period length computed for 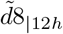 (control) which is identical to the 25.6±0.06h circadian free running period (FRP) period of *S. elongatus* measured at a light intensity of 25 *μmol m*^*−*2^ *s*^*−*1^ ^18^. In *S. elongatus*, nearly all genes have been found to be expressed rhythmically under the control of a circadian (daily) clock. ^19^. Thus, most biochemical processes in cyanobacteria have a natural diurnal periodicity^19,20^. This alludes to a link between exoelectrogenesis and the circadian clock in cyanobacteria. Hypotheses are developed that are consistent with the state of the art model of the cyanobacteria circadian system, putative exoelectrogenic pathways, and the empirical data collected in this work. To give background to the discussion, the circadian system in *S. elongatus* is briefly presented first. In this section, the subscripts |3*h* and |12*h* are dropped for 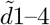 since these modes have the same frequencies in both the 3h:3h and 12h:12h light-dark regimes.

### Overview of the cyanobacteria circadian clock

The circadian system is composed of input pathways, a central biochemical clock and output pathways ^19^. In cyanobacteria, the key input pathways are CikA (circadian input kinase A) and the redox-active protein LdpA (light-dependent period A). The central biochemical clock is the KaiABC protein complex. The output pathway that transmits temporal information from the biochemical clock includes the kinase *Synechococcus* adaptive sensor A (SasA) and the transcription factor regulator of phycobilisome association A (RpaA), which together control gene expression, timing of cell division and other rhythmic biological activity. The KaiABC protein complex keeps time through a highly ordered, sequential cycle of phosphorylation by ATP (day) and dephosphorylation to ADP (night) over a diurnal period ^19^. The cycle is independent of the cell division rate. The biochemical clock identifies day and night *via* the redox state of the plastoquinone pool in the photosynthetic electron transport chain. During the day, the plastoquinone pool is reduced by electrons from upstream photosystem II (PSII), but is rapidly oxidised when light is switched off. CikA and KaiA can only bind to oxidised quinones, and in turn, CikA interacts to KaiC only when bound to an oxidised quinone, signalling the onset of night. Secondly, KaiC senses changes in the ATP/(ATP + ADP) ratio which decreases gradually during the dark interval (irrespective of when the light goes off), signalling the length of night in a light-dark period. The second input component, LdpA, is an iron-sulfur centre redox-active protein that senses changes to the electron transport chain that are dependent on light intensity. Temporal information from the biochemical clock is relayed to gene expression *via* SasA/RdpA ^19,21^. During the day, SasA binds to KaiC promoting SasA auto-phosphorylation followed by phosphotransfer to RdpA to form RdpA-P. RdpA-P is the active form of RdpA and is accumulated over the course of the day, peaking at the day-night transition (dusk). RdpA-P binds directly to about 100 targets in the *S. elongatus* genome, activating dusk-peaking class 1 genes, while repressing the formation of dawn-peaking class 2 genes. Over the course of the night, CikA removes the phosphoryl groups from RdpA-P. In addition to RpaA, there is evidence that the transcription regulator RpaB, which regulates gene expression in response to environmental stress conditions such as photo, thermal or oxidative stress, also plays a part in the output pathway by working cooperatively with RpaA. RpaB can inhibit the phosphorylation of RpaA ^22^. Oscillations in the phosphorylation of RpaB are however only present when cells are grown under a 12h:12h light-dark period ^19^.

### Three modes are identified as candidate circadian rhythms

To differentiate which of the modes may be under circadian clock control, the definition of a circadian rhythm is used. A circadian rhythm is (1) persistent; (2) temperature compensated; and (3) resettable following a dark pulse ^19,20^. The first criteria eliminates the impersistent modes that are primarily active at dawn, 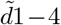 and 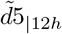. The second criteria does not apply here since all experiments were run under constant temperature. The third criteria rules out 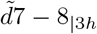 since these did not reset after a dark pulse. This leaves modes 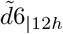 and the analogous pairs 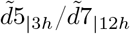, and 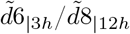 which are persistent and reset after a dark pulse, as candidate rhythms under circadian clock control.

### Inverse of the instantaneous frequency is free running period, while peak-to-peak duration is entrained to light-dark period

It is suggested that the 25.8h median period computed from the inverse of the median instantaneous frequency for mode 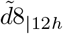 reflects the free running period of *S. elongatus* circadian rhythms. The shorter 24h peak-to-peak duration results from a reset of the oscillation initiated by the dark pulse after 12h of light. The dark pulse interrupts the natural trajectory of the oscillation and entrains the peak-to-peak duration to the light-dark period.

### Shorter peak-to-peak durations and median periods under iron stress are thought to be due to retrenchment of LdpA

Mode 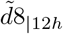 was found to oscillate with median periods that are 1.2–1.4h shorter than the control in the Fe (-)|Air|12h and Fe (-)|20% CO_2_|12h conditions. Further the mean peak-to-peak duration was 1h shorter than the control in the Fe (-)|Air|12h condition. Similarly, the duration between the main dusk-timed peak and lower nighttime peak in 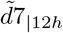 was found to be *≈*1.2h shorter under iron limitation. During culture growth, Fe(-)|Air and Fe(-)|20% CO_2_ cells exhibited a significant drop in chlorophyll *a* content, as well as substantial blue shifts in absorption spectra over time, particularly the 633 nm (phycocyanin) and 683 nm (chlorophyll *a*) absorption peaks, Figs. S3. The pigment changes were more severe in the presence of 20% CO_2_. These are well-documented characteristic responses to iron starvation in *S. elongatus* that are part of a broader rearrangement of the photosynthetic system, including reducing the amount of iron/iron-sulfur rich photosynthetic proteins ^18,23–25^. Interestingly, mutant cells lacking the gene encoding the LdpA protein exhibit these same phenotypic changes, namely: oscillatory periods that are one hour shorter due to insensitivity to light gradients (*i*.*e*., the circadian period length does not increase with reducing light intensity as per Ashcoff’s rule); and reduced phycocyanin content ^18,19^. Since LdpA also contains an iron-sulfur core, it is suggested that the shorter 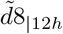 and 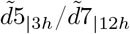 periods observed in the iron depleted conditions are the result of retrenchment of the protein as part of the ensemble of adaptations to reduce iron demand.

### The FM-AM mode 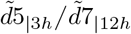 thought to be linked to *psbAI* expression rhythm

The analogous pair 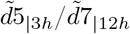 oscillates with three peaks per period, the first in the early day, the highest peak just before dusk, and the final lowest peak in the nighttime *≈*8.68h after the second (control). Strikingly, the timings and amplitude ratios of the second and third peaks are reminiscent of the expression rhythm of the photosynthesis gene *psbAI*. The gene encodes for the D1 reaction centre protein of PSII, in particular the D1:1 form, that plays a key role in the initiation of photosynthesis. At a light intensity of 15 *μmol m*^*−*2^ *s*^*−*1^, the gene is expressed with two peaks within a diel period: a higher dusk-timed peak and a lower nighttime peak ca. nine hours later ^26^. The parallels hint that the biochemical pathway responsible for 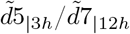 oscillations are linked to the D1:1 protein and by extension, the PSII complex. Furthermore, it is known that under adverse conditions, *S. elongatus* cells preferentially express D1:2, the stress induced form of D1 encoded by *psbAII* and *psbAIII* genes, and repress expression of *psbAI* ^27^. This would explain the diminishing amplitude of the highest, dusk-timed, 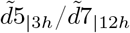 peak observed as nutrients in the media deplete over time. While the hypothesis requires further investigation, that the mode embedding the largest *variable* energy of the photoresponse (in non-stressed conditions) is directly linked to the PSII complex is consistent with putative exoelectrogenic pathways, if not an intuitive conclusion.

### The two modes exclusive to diel regimes speculated to be linked to RpaB regulated genes

Mode 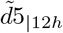 and 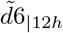 were found to be exclusive to a 12h:12h light-dark period, with 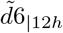 exhibiting its highest peaks at dawn across all operating conditions. The result hints that the biochemical process underlying the oscillations are linked to proteins regulated by RpaB in response to photo-stress, since RpaB phosphorylation only occurs under diel periods. It should be noted that RpaB regulates both circadian controlled genes (*via* RpaA) and arrhythmic genes (independently of the circadian system) ^19,22^. Thus, while 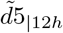 was ruled out as a circadian rhythm on account of its impersistence, it is plausible that it is regulated by RpaB.

### Mode 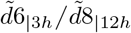 speculated to be linked to intracellular glycogen levels

The analogous pair 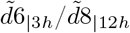 was the only mode to exhibit an increase in amplitude over time as media concentration depleted (control). Further, the amplitude of the mode is uniquely higher in the iron depleted conditions. The implication then, is that the biochemical process responsible for 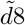 is linked to an iron induced accumulation of a metabolite or the over-expression of a protein that participates in the exoelectrogenic pathway. Of the suite of iron starvation induced adaptations in *S. elongatus*, increase in flavodoxin, expression of iron starvation induced outer membrane proteins, and increase in glycogen content are the most relevant suspects consistent with putative exoelectrogenic pathways.

To reduce iron demand in iron depleted conditions, levels of the redox protein flavodoxin increase in replacement of the retrenched electron mediator ferrodoxin ^23^. However, since flavodoxin, like ferrodoxin, is an intracellular electron carrier yet to be implicated in electron export, there no evidence to support that replacement of one with the other alters exoelectrogenic rates. Furthermore the expression rhythm of *isiB*, the gene encoding flavodoxin, peaks in the early day just after dawn, which is inconsistent with the late day peak of 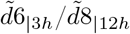 (control) ^28^.

Three outer-membrane proteins of 92, 35 and 48-50 kDA mass are induced in iron starved *S. elongatus* that are otherwise unexpressed in cells grown in iron replete media ^29^. Further, as highlighted in the introduction, cyclic voltammetry experiments in the research group provided evidence of distinct outer membrane redox activity in iron starved cells ^7^. Taken together, the evidence supports the notion that 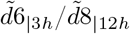 oscillations may be linked to one or more of the three iron starvation induced proteins. However, little is know of the nature of the proteins and there is no evidence for or against the view that the proteins are expressed rhythmically. To the contrary, given that the proteins are expressed in response to stress, it is more likely that they are under environmental regulation.

The most compelling hypothesis is that 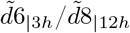 is linked to fluctuations in cellular glycogen levels. *S. elongatus* intracellular glycogen levels oscillate under circadian control, increasing over the course of the day and peaking at dusk (in standard laboratory conditions) ^30,31^. In parallel, current output has been reported to be proportional to intracellular glycogen content in *Synechococcus* sp. (strain not declared) BPVs operated in the presence on 3% v/v CO_2_ ^32^. Finally, glycogen levels in *S. elongatus* increase under iron limitation, possibly to reduce the need for CO_2_ fixation and thus stress on the retrenched photosynthetic system ^33^. Combined, the studies offer a coherent model for a glycogen-coupled electron flux, with a circadian rhythm, that peaks near dusk, and is amplified in iron limited conditions.

### Phase shifts under iron stress and in the presence of CO_2_ hypothesised to be due to changes to intracellular ATP/(ATP + ADP) ratio

*In vitro* and *in vivo* experiments have shown that when the intracellular ATP/(ATP + ADP) ratio (%ATP henceforth) is reduced below normal (typically *≈*80% in the light, falling to *≈*50% within 2–3h of darkness), circadian rhythms advance towards dawn ^30,34^. The more extreme the reduction in %ATP during night, the greater the phase advance after a dark interval. Thus, the advancement of modes 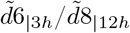 and 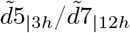 (early day) peaks towards dawn in the Fe (-)|Air condition is hypothesised to be caused by lower intracellular %ATP in the cells at night. This is consistent with lower photosynthetic activity, ergo reduced accumulation of ATP, during the day under iron limitation.

The phase shifts observed in the Fe (-)|20% CO_2_ modes were more extreme. There was evidence that the presence of CO_2_ adversely affected the functioning of the CikA protein that is responsible for cleaving the phosphorous group from Rpa-P during the night. Fe (-)|20% CO_2_ cells, were found to be elongated by up to 1.6 *μm* relative to Fe (-)|Air cells (Fig. S5) and by *≈*0.5–0.8 *μm* relative to cells grown in standard conditions (reported in previous work) ^16^. In *S. elongatus*, this phenotype indicates elevated levels of RpaA-P, which implies a reduction/interruption in normal CikA activity ^19,22^. Fe (-)|Air cells exhibited the opposite phenotype, with shortened cells that were only 62–80% the length of iron replete cells (Fig. S5). Cells lacking CikA overaccumulate glycogen during the day, and exhibit higher %ATP than wild-type cells over the course of night^30^. Consequently, the mutant cells exhibit reduced ability to reset phase following a dark pulse. It is therefore speculated that the elevated levels of CO_2_ increase nighttime intracellular %ATP thus causing the peculiar phasing of oscillations observed in the Fe (-)|20% CO_2_ modes.

### The highest frequency modes speculated to be linked to ephemeral photosynthetic electron transport chain activity upon illumination

It was demonstrated in the results that impersistent modes 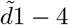 are ferric sensitive, *i*.*e*., their amplitudes decay and recover in response to ferric ion concentration. The behaviour correlates to the pigment changes under iron starvation discussed above, and that are known to be reversible upon iron replenishment ^24,33^. Taken together with the observation that the modes are ephemerally active when there is a light transition, it is is speculated that the oscillations in 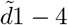 are due to short-lived phenomena linked to the activation/deactivation of the photosynthetic electron transport chain immediately after illumination/darkness.

### The probable genesis of four modes remains unclear at this time

At this time, it is not clear to the authors, based on the circadian model and putative exoelectrogenic pathways, what the likely genesis of modes 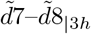 might be and why the modes only appear under the 3h:3h light-dark regime. Similarly, while 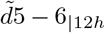 are thought to be under RpaB regulation, it is unclear at this time what the probably genesis of the modes are.

### Combining ICEEMDAN with machine learning is a promising approach to modelling photobioelectrochemical systems

One of the key motivations for this work was to decompose BPV current density into more regular, physically meaningful subcomponents that can be modelled with machine learning or other modelling approaches. It is evident that ICEEMDAN achieves this objective. Each current density profile is decomposed into a maximum of eight modes that are predictable and, individually, substantially easier to model than the parent observed current density. Mode 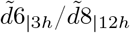 for example, is easily modelled with a simple sin/cosine function. As the physical meanings of the modes are unravelled, for example if the proposed link between ferric ion concentration and the attenuation of amplitude or that between intracellular %ATP and phase shifts are confirmed, simple but robust models can be developed that take as input forecasts of the environmental nutrient and intracellular metabolite concentrations to model individual IMFs, which can then be summed to provide a prediction of BPV performance over time. A combination of empirical mode decomposition and LSTM networks has been recently used to successfully model complex signals in nuclear power plants ^35^.

Tab. 1 summarises the results and hypothesis presented and discussed.

**Table 1.**
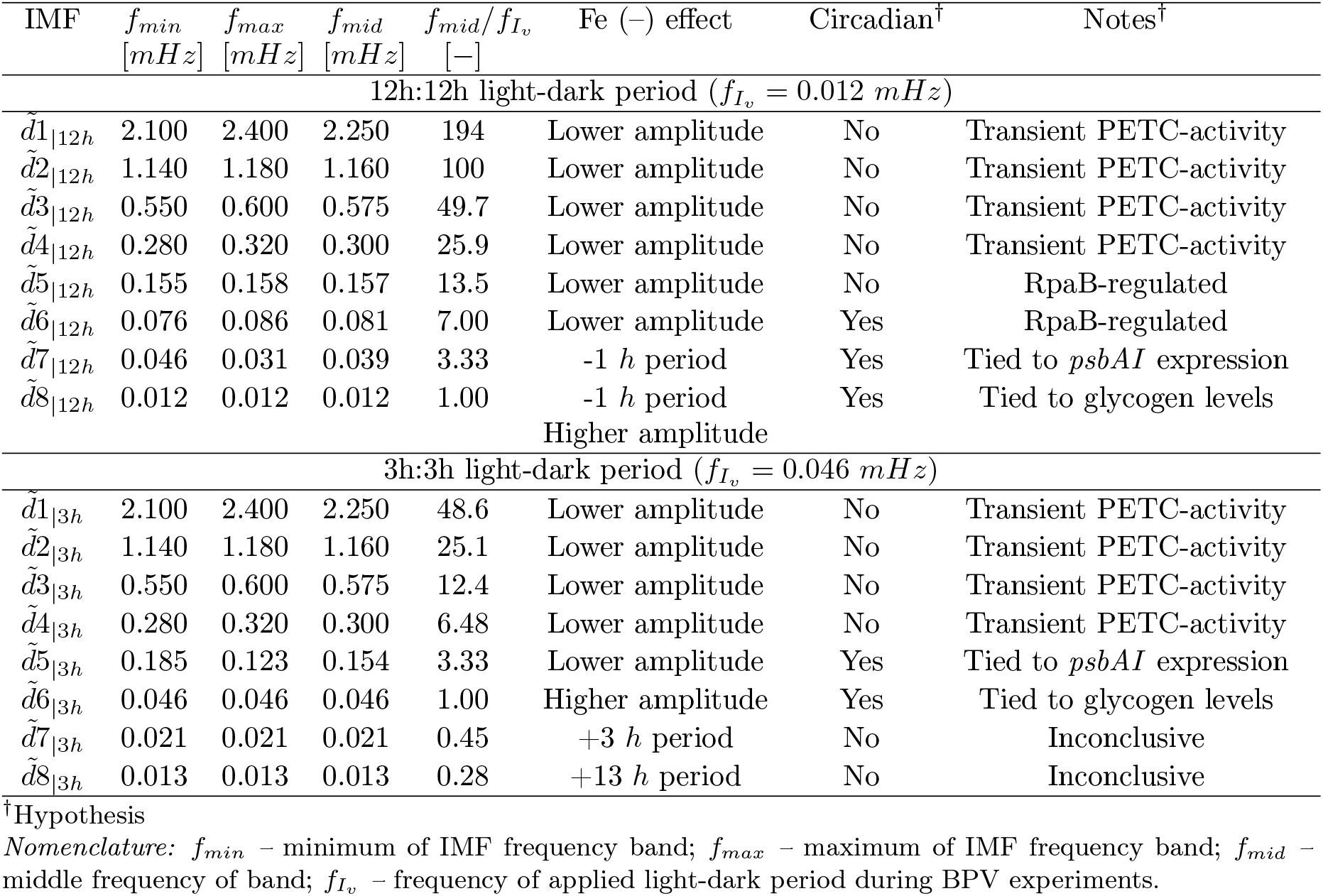
Summary of frequency bands obtained *via* the Hilbert-Huang transform and hypotheses for their physical meaning. The *Fe (–) effect* column shows the response to iron-depleted conditions. In addition, all IMFs exhibited some form of a phase shift under iron-depleted conditions, which was made more severed in a 20% CO_2_ atmosphere.

## 5 Conclusions

Ten characteristic oscillation modes making up *S. elongatus* exoelectrogenic signals have been identified, two of which only exists under 3h:3h light-dark periods, and two of which only exists under 12h:2h light-dark periods. Of these ten, three modes are thought to be under circadian control. Amplitude differences and phase shifts across different operating conditions in these modes are consistent with the prevailing circadian model of the cyanobacterium. Hypotheses have been laid out for the physical meaning of six modes, while four remain unexplained at this time. To test the hypotheses and further interrogate the physical meaning of the modes, the experiments and analysis conducted in this work should be repeated with mutant cells genetically altered to test the proposed hypothesis (*e*.*g*., Δ*psbAI* mutants to test the hypothesis for the genesis of mode 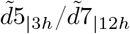, Δ*ldpA* mutants to test the hypothesis for the shortening of period 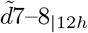 under iron starvation *etc*.). The experiments and analysis should also be repeated in the presence of site specific respiratory and photosynthetic electron transport chain inhibitors which we expect will affect modes 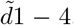. Finally, the experiments and analysis should be repeated under constant light to test the discrepancy between the median period (calculated from the median instantaneous frequency) and the peak-to-peak duration of mode 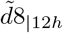. Circadian influence in exoelectrogenesis as proposed in this work may offer an explanation for the difference in patterns (amplitudes, timings of peaks and troughs *etc*.) of exoelectrogenic signals across cyanobacteria species. For example, *Synechocystis* is known to have lower amplitude and less accurate circadian rhythms than *S. elongatus* ^28^. This is consistent with higher peaks and troughs reported in *Synechococcus* chronoamperograms ^8^. Finally, a confirmation of the hypotheses developed in this work may offer a bioelectrochemical approach for performing simple chronobiology experiments that eliminates the need for engineering reporter strains and extensive fluorescence measurements typical in the study of cyanobacteria circadian rhythms, while increasing the time resolution of experiments.

## 6 Methods

### 6.1 Culturing and characterisation

All cultures were grown at 30^°^*C*, a white light intensity of 21.0±0.3 *μmol m*^*−*2^ *s*^*−*1^ and a shaking speed of 120 *rpm*.

#### Stock culture

A stock culture of *Synechococcus elongatus* sp. PCC7942 (Pasteur Culture Collection) was grown in liquid blue-green BG11 media ^36^. The media was replenished when the stock culture entered the decline phase. The stock culture was tested for axenicity prior to starting the experimental cultures by plating on tryptic soy plates and leaving in the dark to ensure no growth.

#### Experimental cultures

Experimental cultures were started by re-suspending biomass pellets obtained from the stock culture in exponential phase by rapid centrifugation (4000 *×* g for 10 minutes) in fresh media following two washing steps. The cultures were grown for 4 days (96 *h*), before inoculating in the BPV devices.

For the iron deplete cultures, Fe (-)|Air and Fe (-)|20% CO_2_, the BG11 medium was modified by replacing ammonium ferric citrate by an equal molar amount of ammonium citrate ^7^. In addition, the medium for the Fe (-)|20% CO_2_ cultures was buffered to pH 7.0 using HEPES-NaOH to prevent excessive acidification of the culture by CO_2_. The 20% CO_2_ atmosphere was induced by growing the cultures in Erlenmeyer flasks with two port caps. The inlet was supplied by a 20% v/v CO_2_ gas supply at 10 *ml min*^*−*1^. Fig. S7 shows the P&ID of the experimental set-up. The inlet and outlet ports were fitted with 0.2 *μm* filters to maintain a sterile environment within the flasks.

All glassware used to prepare media and grow the iron depleted cultures were soaked in nitric acid overnight to reduce traces of solids and therefore iron to a minimum ^7^. All media were prepared with Millipore^®^ ultra pure reverse osmosis water.

#### Determination of cell number

Cell number, N (*cells ml*^*−*1^), was estimated from optical density at 750 nm (OD750). To account for the effects of cell size and medium composition on optical density readings, independent standard curves for converting OD750 readings to N were calibrated for each growth condition ^37^. To generate the curves, OD750 measured using a Thermo Scientific Evolution 201 UV–Visible spectrophotometer, and N measured using a Beckman Coulter^™^ Z2 particle counter, were recorded over 216 hours (Fig. S1a–c). A second order polynomial was then fit to the data to produce equations for estimating N from OD750 (Fig. S1d–f and Tab. S1).

#### Determination of chlorophyll *a* content and cell size

Chlorophyll *a* content and cell size were determined as described in previous work ^16^.

### 6.2 BPV device and operation

BPVs were operated at 30^°^*C* and a white light intensity of 24±0.3 *μmol m*^*−*2^ *s*^*−*1^.

#### BPV architecture

BPV architecture and assembly was as described in recent work ^12^. To recap, the membrane electrode assembly (MEA) was made up of a porous Toray carbon paper anode, a nitrocellulose membrane (0.22 *μm* pores), and an Alfa Aesar platinum coated carbon paper cathode with 3 *mg m*^*−*2^ Pt loading. The growth medium served as the electrolyte and no exogenous mediators were used.

For the Fe (-)|20% CO_2_ condition, a 20% CO_2_ atmosphere was induced by sealing the anode chamber with a two port cap, and supplying the inlet with 10 *ml min*^*−*1^ of a 20% v/v CO_2_ gas supply. The inlet and outlet were fitted with 0.2 *μm* filters to maintain a sterile environment (Fig. S7–8).

#### Operation

BPVs were inoculated with 5 ml of culture started from biomass pellets harvested from the experimental cultures (4000 *×* g for 10 minutes) after four days of growth. The pellets were resuspended in fresh medium to a cell concentration of 6.78*×*10^8^ *cells ml*^*−*1^ after one washing step. BPVs were left to stand at open circuit for four days under a 12h:12h light-dark cycle to allow cells to colonise the carbon anodes and form a biofilm (Fig. S9a). Following biofilm formation, the operating procedure shown in Tab. 2 was followed. Voltage measurements were taken every two minutes and converted to current density *J* using Ohm’s law,

**Table 2.** Operating procedure for BPVs.

|  |  |
| --- | --- |
| 1. | Open circuit potential (OCP) measured. |
| 2. | 33 $M\Omega$ external resistor connected; light-dark period set to 3h:3h. |
| 3. | Voltage allowed to stabilise; repeatable photoresponse recorded. |
| 4. | Polarisation curve measured in the dark<sup>‡</sup>. |
| 5. | 33 $M\Omega$ external resistor connected; light-dark period set to 12h:12h. |
| 6. | Voltage allowed to stabilise; repeatable photoresponse recorded. |
| 7. | Polarisation curve measured under illumination. |
| 8. | Media refreshed; 33 $M\Omega$ external resistor connected; light-dark period set to 3h:3h<sup>‡</sup>. |
| 9. | Voltage allowed to stabilise; repeatable photoresponse recorded<sup>‡</sup>. |

<sup>‡</sup> Excluding Fe (-)|Air BPVs.

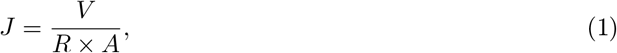

where *V* is the measured voltage, *R* the applied external load, and *A* the electrode geometric area.

### 6.3 Statistical analysis

Cultures were grown in three independent replicates for each growth condition and three independent BPVs were run for each growth condition. All readings were conducted for each replicate and means taken. For confocal image processing, the weighted average means and standard deviations were calculated to account for the different number of cells in the image of each sample. Student’s t tests at 5% significance level was used to test the difference in cell sizes.

### 6.4 Hilbert-Huang transform

#### Hilbert transform

The Hilbert transform enables the identification of the instantaneous frequency and energy of a signal by creating an analytic signal (a complex-valued function) from a measured time series.

For a time series *X*(*t*), its Hilbert transform *Y* (*t*) is given by Eq. 2,

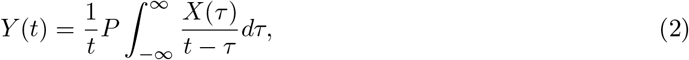

where P is the Cauchy principal value. The signal *X*(*t*) and its Hilbert transform *Y* (*t*), both real, are then paired to form the analytic signal *Z*(*t*). The analytic signal is a complex number with *X*(*t*) as the real part and *Y* (*t*) the imaginary part. The analytic signal, *Z*(*t*), can then be represented in polar form using Euler’s formula, Eq. 3,

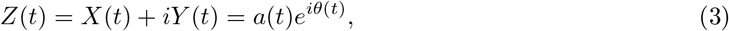

where *i* is the imaginary unit. The modulus of the complex number, *a*(*t*), is defined as the instantaneous energy and is given by

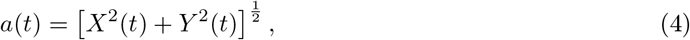

while the argument of the complex number, *θ*(*t*), is defined as the instantaneous phase of *X*(*t*) and is given by

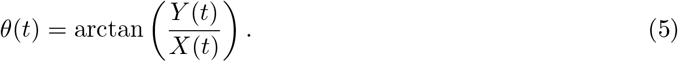

Finally, the instantaneous frequency of *X*(*t*) is defined as derivative of the instantaneous phase, *ω*(*t*),

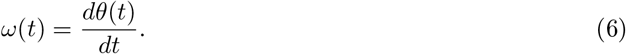

The maximum frequency that can be extracted is limited by Δ*t*,

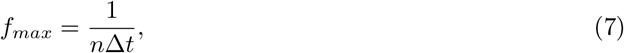

where *n* is the minimum number Δ*t* required to define a frequency. In the original implementation of HHT, it was determined that at least five data points are required to define a frequency, translating to *n* = 4. In this work, Δ*t*=120 *s* giving a maximum extractable frequency of 2.1 *mHz*. However, to interpret *ω*(*t*) as the instantaneous frequency, some restrictions apply. First, given that Eq. 6 is a single value function, *i*.*e*., there is only one frequency *ω* at a given time t, the data to be transformed must also represent only one component at the time t ^13^. Physically, this may be conceptualised as the oscillating data having the same number of zero mean crossings and extrema (maxima, minima) per unit time. Secondly, the data must be symmetric around the *local zero mean* level, otherwise Eq. 6 may result in physically meaningless negative frequencies ^13^. The Empirical Mode Decomposition (EMD) was developed to decompose nonlinear and/or non-stationary signals into components that meet the above restrictions.

#### Empirical Mode Decomposition (EMD)

The goal of EMD is to decompose a signal that contains multiple dynamic oscillations superimposed on each other, into a finite set of physically meaningful components ^13,35^. One key attribute of EMD is the definition of the *local mean* of a signal as the mean of the upper and lower envelopes determined from its local extrema. That is, the separate lines connecting the local maxima [upper envelope] and local minima [lower envelope] by a smooth interpolation over the timespan of the signal.

With the local mean as the reference point for oscillations, high frequency oscillations (oscillations around the local mean) can be extracted from the low frequency oscillations (embedded within the local mean) in an iterative sifting process. The result is a series of components, from high to low frequency, called intrinsic mode functions (IMFs), each encapsulating oscillations of a characteristic timescale (time lapse between successive extrema) intrinsic to the data. For a particular IMF, each cycle (between zero crossings) contains only one mode of oscillation, *i*.*e*., with no superimposed riding waves ^13^.

The formal definition of an IMF as given in the original implementation of EMD, is a function that satisfies two conditions ^13^:

1. The number of extrema and the number of zero crossings are equal or differ at most by one.
2. The mean value of the envelope defined by the local maxima and the envelope defined by the local minima is zero at all points.

The first condition limits each IMF to one component at any point in time (no riding waves), allowing calculation of instantaneous frequency using Eq. 6. The second condition ensures symmetry around the local mean to prevent negative frequency values. Algorithm 1 shows the EMD pseudo-code.

##### Algorithm 1

EMD

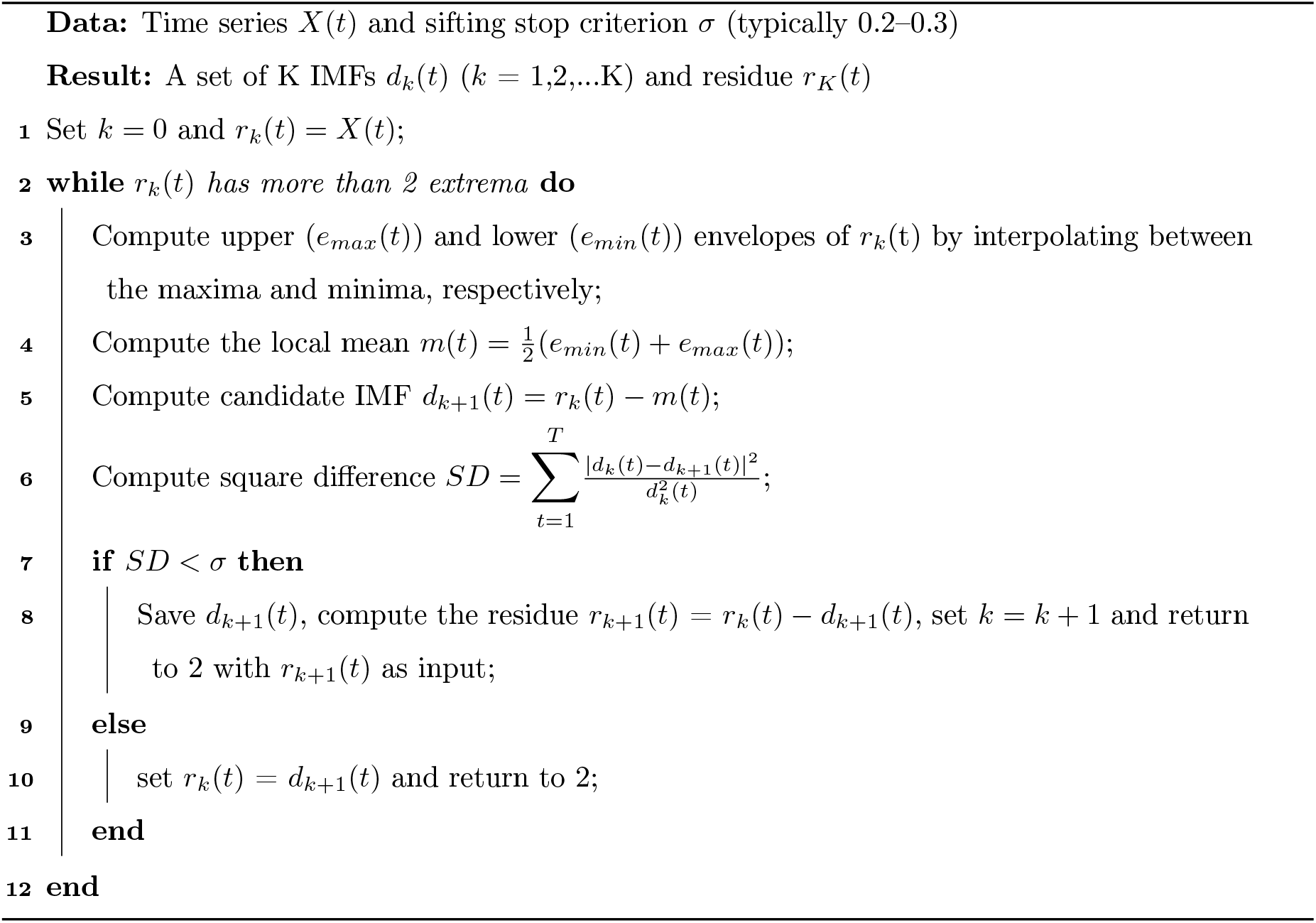

The original signal can be reconstructed as the sum of the IMFs and the final residue which is a monotonic function,

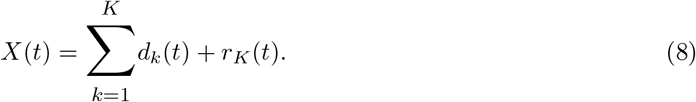

It should be noted that since the local mean changes over the course of the signal, what is considered a high frequency in one part of the signal may be considered low frequency in another part of the signal. Therefore, both the frequency and the amplitude of the resultant IMFs are allowed to vary with time (AM-FM), adapting to the local characteristics of the signal. A caveat of this adaptability is that one IMF may include oscillations with vastly differing timescales, or oscillations of similar timescales may occur in different IMFs ^14^. This is termed ‘mode mixing’. To reduce mode mixing, several improvements to the EMD have been proposed, including the Ensemble EMD (EEMD), Complete EEMD with Adaptive Noise (CEEMDAN), and most recently the Improved CEEMDAN (ICEEMDAN) ^14,38,39^. In this work, ICEEMDAN is used to obtain similar timescales in each IMF.

#### Improved Complete Ensemble EMD with Adaptive Noise

To reduce mode mixing, the ICEEMDAN approach creates copies of the residue *r*_*k*_ at each stage of decomposition and adds the *k*th IMF of different realisations of controlled white Gaussian noise to each copy, resulting in an ensemble of noisy residues ^14^. Then, the local mean of the residue is taken to be the average of the local means of its noisy copies. This helps improve the estimation of the local mean, resulting in IMFs of similar scale ^14^. In algorithm 2, the operator *E*_*k*_(·) produces the *k*th IMF *via* EMD, *M* (·) produces the local mean of the signal it is applied to (as in step 4 in algorithm 1), (·) is the action of averaging over the different copies of residue plus noise, and *std*(·) is the standard deviation.

##### Algorithm 2

ICEEMDAN

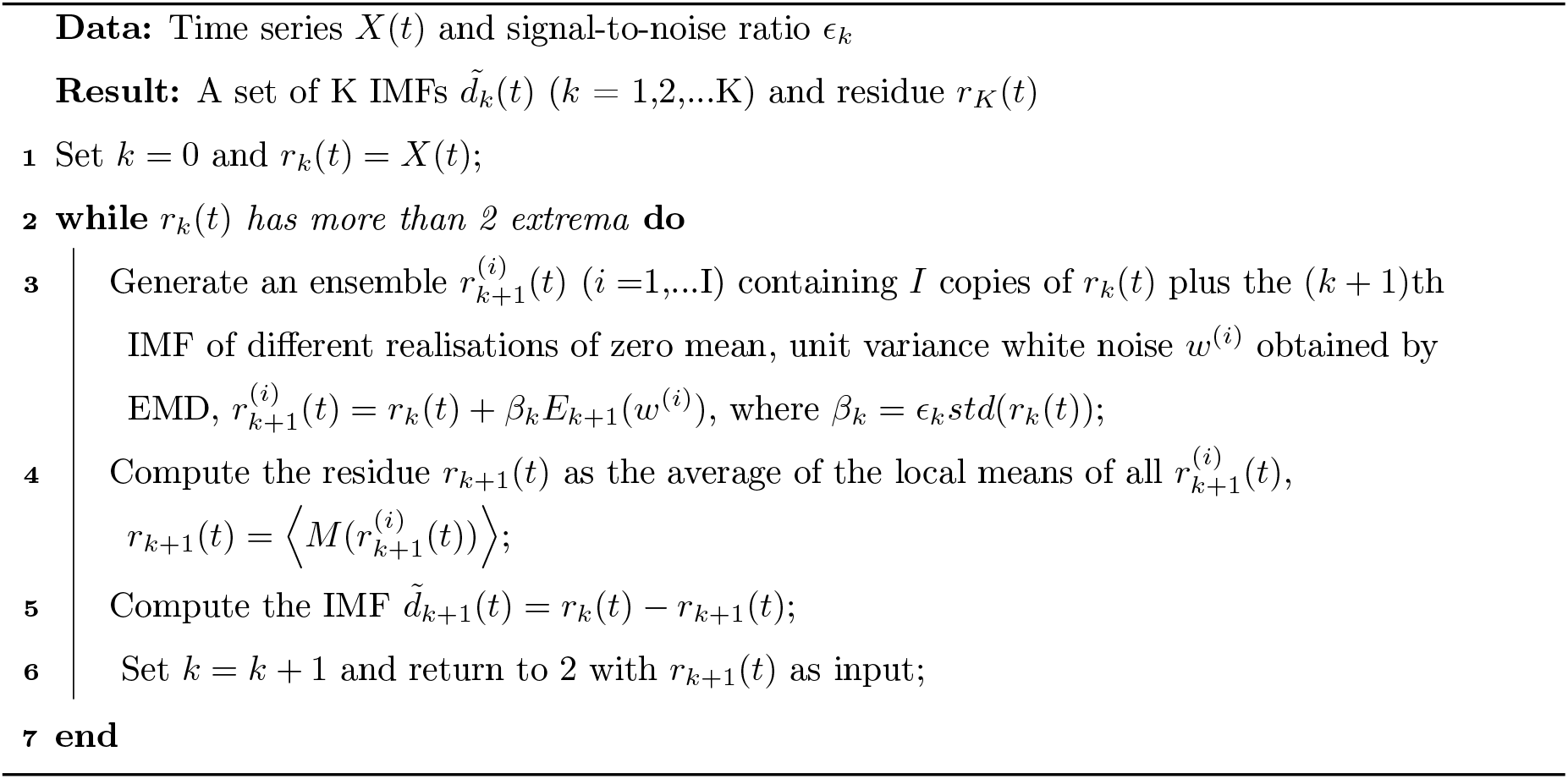

In this work, ICEEMDAN was implemented using the Matlab code developed in the original implementation of the method ^14^.

#### Hilbert spectrum and marginal spectrum

After ICEEMDAN decomposition of *X*(*t*), the Hilbert transform can be applied independently to each IMF 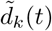, so that the analytic signal in Eq. 3 can be re-written as the sum of the analytic signals of the K IMFs,

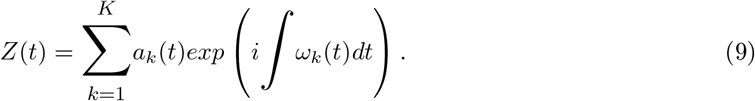

In Eq. 9, the final residue *r*_*K*_(*t*) is intentionally excluded since there is some uncertainty whether the monotonic component is part of a longer oscillation and the instantaneous energy of *r*_*K*_ can be overpowering relative to the energies of 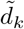^13^.

The information in Eq. 9 can be represented in a three-dimensional diagram called the ‘Hilbert spectrum’, *H*(*ω, t*), with *t* on the x-axis, *ω*_*k*_(*t*) on the y-axis, and *a*_*k*_(*t*) contoured on the frequency-time plane for all K IMFs ^13^.

In addition, the marginal spectrum, *h*(*ω*), is defined as

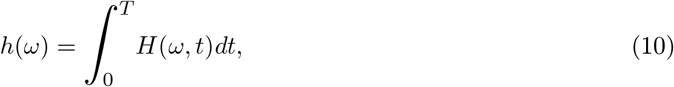

and provides a measure of the total contribution from each frequency value to the variable amplitude.

## Acknowledgements

This work was supported by funding from Cambridge-Africa, Cambridge Trust awarded to T. I. Okedi, and from the British Council Newton Fund Prize [Grant Number RG95201] and Cambridge CARES C4T to Dr A. C. Fisher. We thank Dr Arely Gonzalez and Dr Aazraa Pankan for their BPV operating protocols, device designs and advice. Cyanobacteria icons in Fig. 1 are from biorender.com.

## Author contributions

TIO: conceptualisation, data curation, formal analysis, funding acquisition, investigation, software, validation, visualisation, writing – original draft. KY: funding acquisition, resources, writing – review & editing. ACF: funding acquisition, resources, supervision.

## Materials & Correspondence

Correspondence and material requests should be addressed to TIO and/or ACF.

