## Supplemental information for "Green notes: The rhythms of cyanobacteria exoelectrogenesis as de-composed by the Hilbert-Huang transform"

### $OD_{750}$ to cell number calibration curves

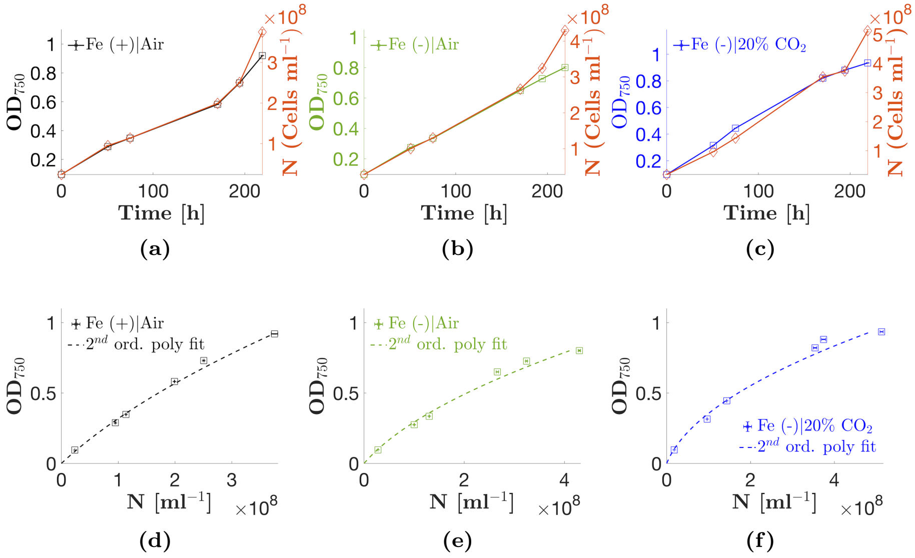

**Figure S.1 – Calibration and standard curves for converting $OD_{750}$ readings to cell number.** **(a)–(c)** show growth curves and cell counts for cultures used to generate the $OD_{750}$-to-cell number (N) calibration curves. The correlation between $OD_{750}$ and N breaks down after ≈192 $h$ (8 $days$) of growth. **(d)–(f)** $2^{nd}$ order polynomial fit standard curves for each growth condition. To generate the fitted curves, the last (216 $h$) $OD_{750}$-N pair from each calibration growth curve is ignored. Error bars on data points show ±1 SEM, n = 5. Equations for the standard curves are shown in Tab. S.1.

**Table S.1** – Equations for the $2^{nd}$ order polynomial standard curves shown in Fig. S.1. The $OD_{750}$ column shows the maximum measured optical density below which the relationship is valid. All cultures were diluted to below the maximum valid $OD_{750}$ and the calculated value N multiplied by the dilution factor to obtain the final cell number.

| **Condition** | $N$ $[cells\ ml^{-1}]$ | $R^2$ | $OD_{750}$ |
|---|---|---|---|
| Fe (+) \| Air | $1.527 \cdot 10^8 \times OD^2 + 2.652 \cdot 10^8 \times OD$ | 0.9988 | $< 0.92$ |
| Fe (-) \| Air | $3.390 \cdot 10^8 \times OD^2 + 2.411 \cdot 10^8 \times OD$ | 0.9851 | $< 0.80$ |
| Fe (-) \| 20% $CO_2$ | $4.414 \cdot 10^8 \times OD^2 + 1.338 \cdot 10^8 \times OD$ | 0.9887 | $< 0.93$ |

### Experimental cultures growth curves

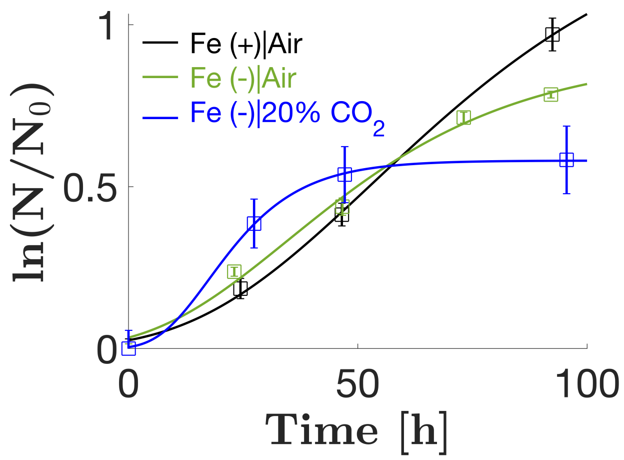

**Figure S.2 – Experimental cultures growth curves.** Specific growth rates ($\mu_{max}$), lag times ($\lambda$) and maximal log of the relative cell number ($A$) were estimated by fitting the Gompertz model to the cell concentration profile [1]. The fitted parameters are shown in Tab. S.2. Cultures were grown from a stock culture at a starting $\mathrm{OD}_{750}$=0.5 and inoculated into BPVs after four days of growth.

**Table S.2** – Gompertz model fitted parameters with 95% CI for the experimental culture growth curves in Fig. S.2

| **Condition** | $\lambda$ $[h]$ | $\mu_{max}$ $[h^{-1}]$ | $A$ $[-]$ | $R^2$ |
|---|---|---|---|---|
| Fe (+) \| Air | 14.8±38.9 | 0.013±0.016 | 1.39±3.69 | 0.9980 |
| Fe (-) \| Air | 5.57±22.4 | 0.011±0.008 | 0.91±0.50 | 0.9919 |
| Fe (-) \| 20% $CO_2$ | 6.39±24.8 | 0.019±0.022 | 0.58±0.08 | 0.9998 |

**Abbreviations:** $\lambda$ - Lag time; $\mu_{max}$ - Maximum growth rate; $A$ - Maximum log of relative cell number, $ln(N_\infty/N_0)$ where $N_\infty$ is the cell concentration at stationary phase.

#### Chlorophyll *a* and absorption peaks profiles

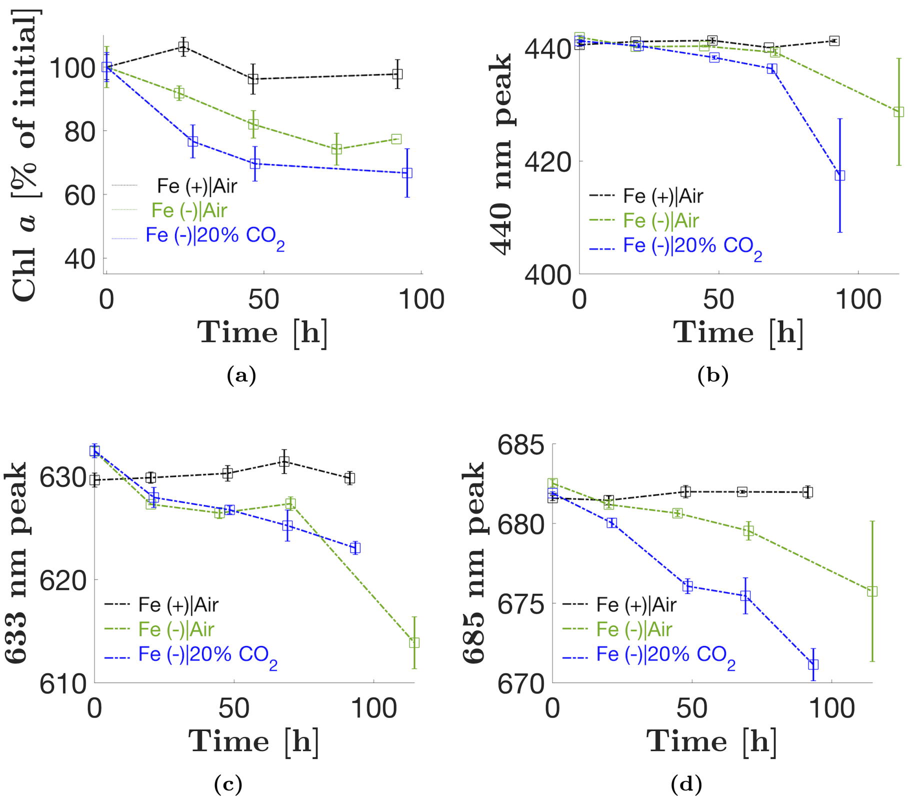

**Figure S.3 – Chlorophyll *a* and absorption peak profiles for each growth condition.** Error bars show ±1 SEM for three independent replicates (n=3). Where error bars are not visible, they are smaller than the marker size.

#### pH profiles

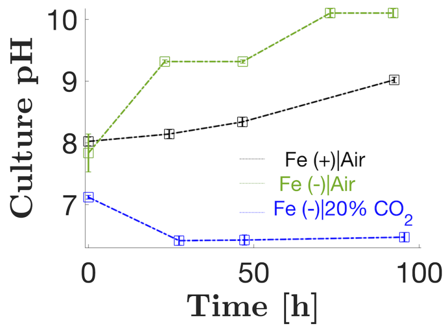

**Figure S.4 – Culture pH profiles**. Error bars show ±1 SEM for three independent replicates (n=3). Where error bars are not visible, they are smaller than the marker size.

### Cell size

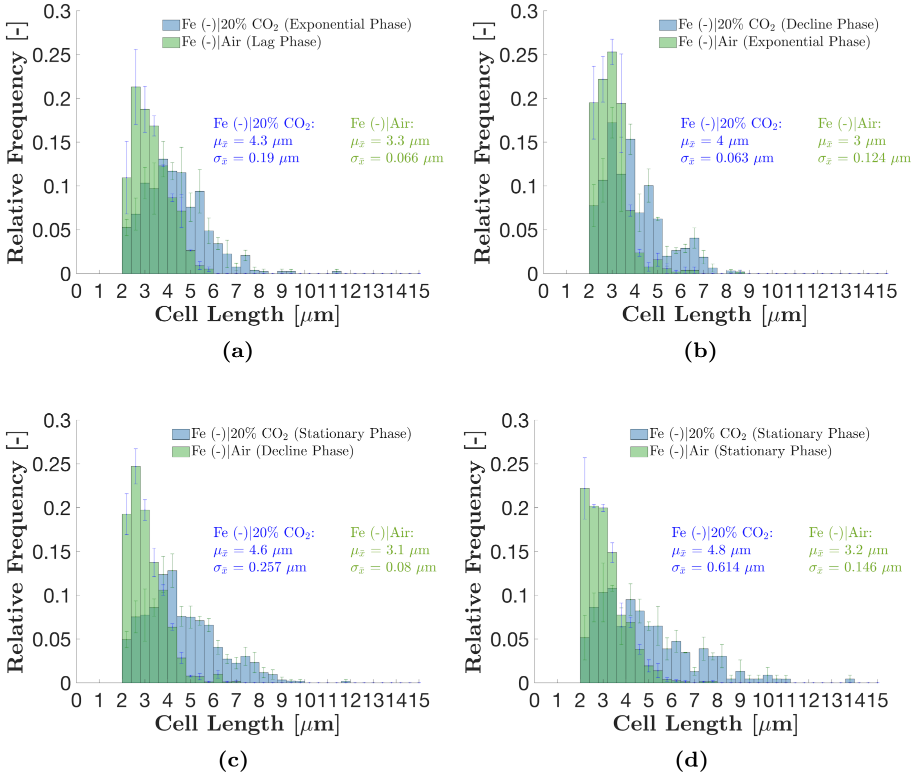

**Figure S.5 – Cell size distribution for Fe(-) cultures. (a)** After 24 hours of growth in media. 3 replicates with n = 133, 92, and 139 cells for the Fe (-) | 20% $CO_2$ cultures. 3 independent replicates with n = 120, 74 and 147 cells for the Fe (-) | Air cultures. **(b)** After 120 hours growth in media**.** 3 replicates with n = 32, 198, and 255 cells for the Fe (-) | 20% $CO_2$ cultures. 3 independent replicates with n = 33, 181 and 166 cells for the Fe (-) | Air cultures. **(c)** After 216 hours of growth in media**.** 3 replicates with n = 188, 103, and 161 cells for the Fe (-) | 20% $CO_2$ cultures. 3 independent replicates with n = 246, 344 and 109 cells for the Fe (-) | Air cultures. **(d)** After 264 hours of growth in media. 2 replicates with n = 117, and 115 cells for the Fe (-) | 20% $CO_2$ cultures. 3 independent replicates with n = 409, 230 and 808 for the Fe (-) | Air cultures. Error bars show ± 1 SEM. All averages and standard deviations are weighted by sample size.

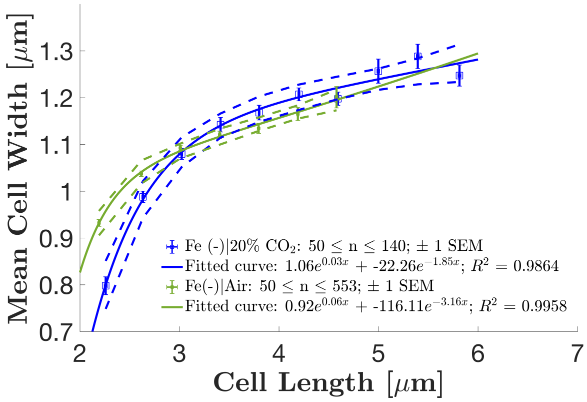

**Figure S.6 – Cell width vs. cell length**. The plot was created using data collected over the duration of the experiment from 2,526 and 1,169 cells from the Fe(-)|Air and Fe(-)|20% $CO_2$ conditions respectively. The data were discretised by cell length into bins of 0.4 $\mu m$ wide. Mean values were calculated from bins with at least 50 data points to produce the plot. Dotted lines are 95 $\%$ confidence bounds.

### $CO_2$ experiments P&ID set-up

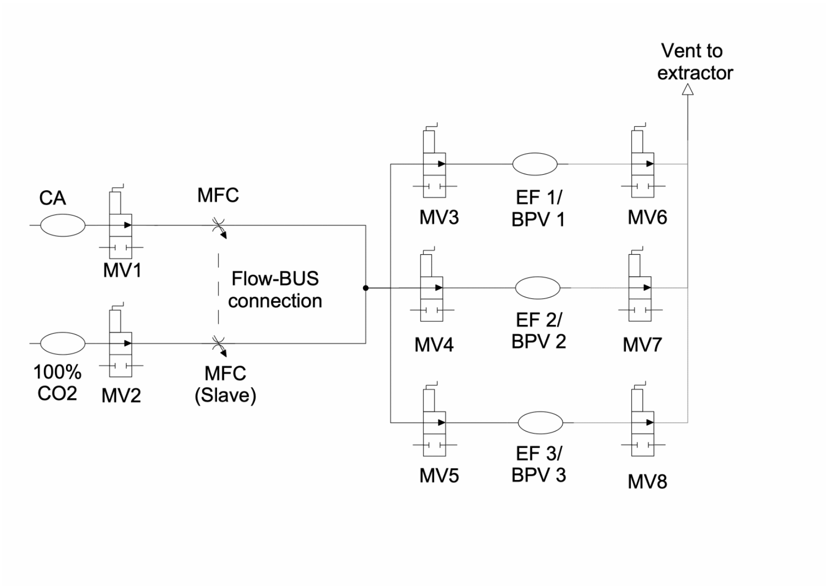

**Figure S.7 – P&ID of set-up for $CO_2$ experiments**. Compressed air (8 $ml\ min^{-1}$) and 100% $CO_2$ (BOC, 2 $ml\ min^{-1}$, ) were mixed into a single stream (10 $ml\ min^{-1}$) with a 20% $CO_2$ concentration. The ratio of flow rates of the two streams was regulated by mass flow controllers connected in a FLOW-BUS network. The controller regulating $CO_2$ flow was a slave to the controller regulating compressed air flow (set-point of the $CO_2$ controller determined from measured compressed air flow rate using a user defined 4:1 Air:$CO_2$ ratio). This ensured that the desired $CO_2$ concentration remained constant in the input ports regardless of any fluctuations in the compressed air flow. The outlet port was vented to an extractor. Abbreviations: CA - Compressed air; MV - Manual valve; MFC - Mass flow controller (Bronkhorst EL-FLOW); EF - Erlenmeyer Flask; BPV - Biophotovoltaic device.

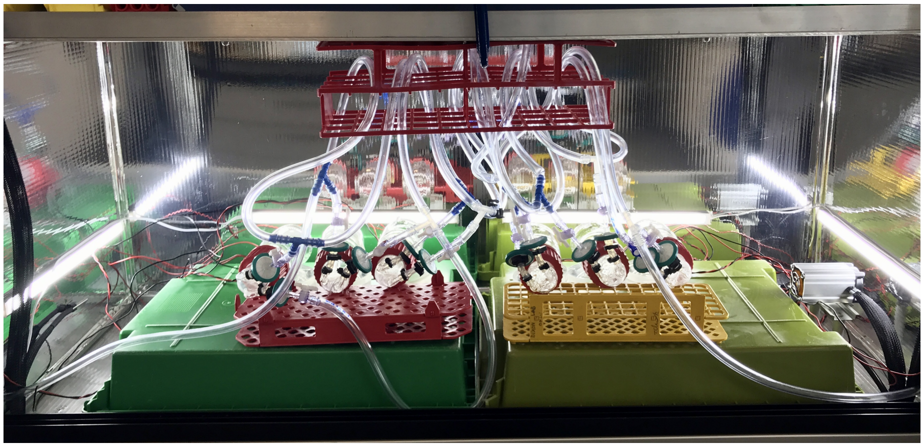

**Figure S.8 – BPV set-up in experimental rig**. Device architecture is as previously reported [2]. Connections are as shown in Fig. S.7.

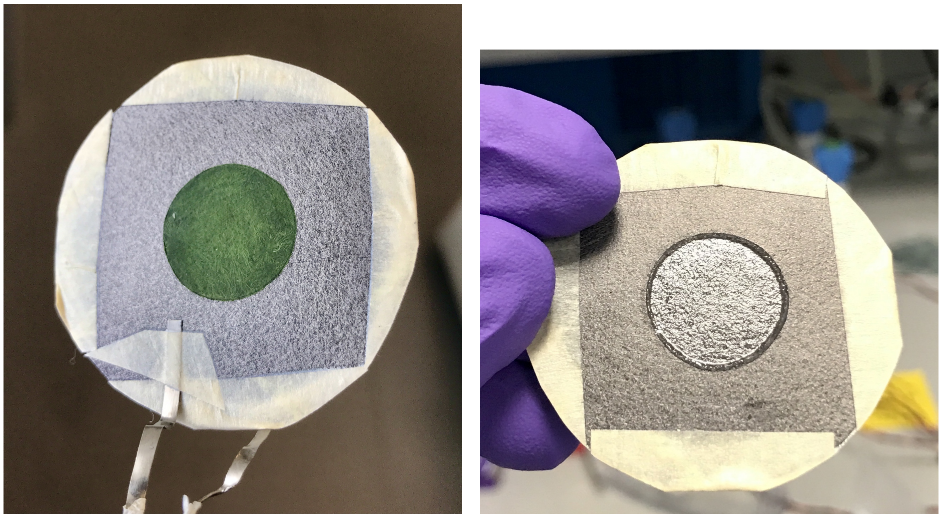

(a) (b)

**Figure S.9 – Biofilm on carbon anode**. **(a)** *S. elongatus* biofilm on carbon anode at the end of the experiment (315h) from a Fe(+)|Air device. **(b)** Anode from Fe(+)|Air media only device at the end of experiment showing precipitation of media salts.

### BPV polarisation curves

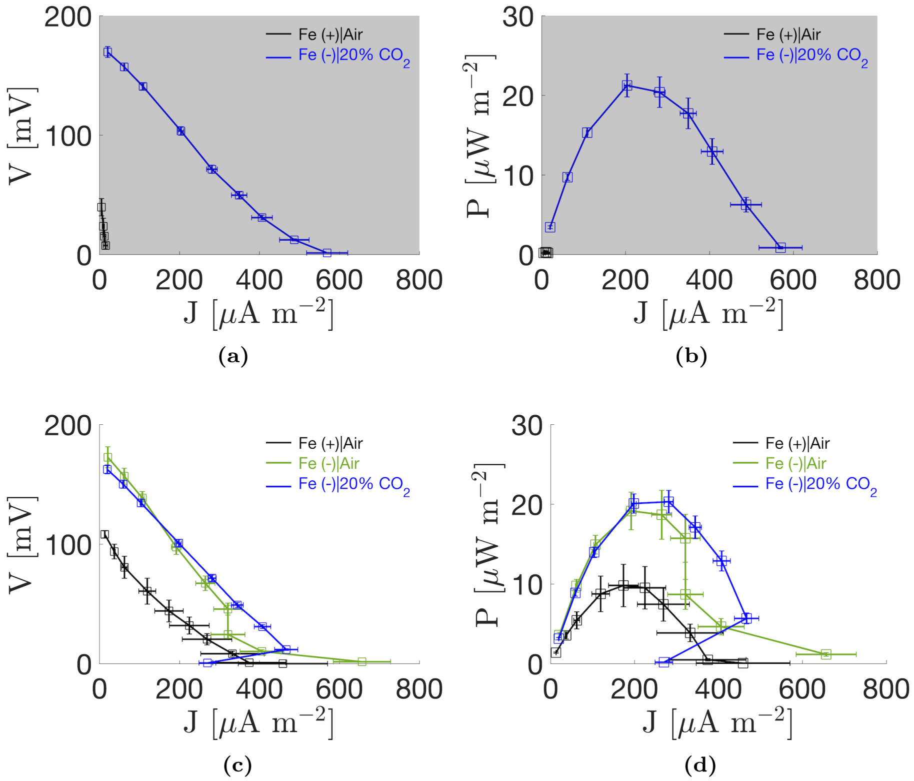

**Figure S.10 – BPV polarisation curves for each growth condition**. **(a)** Polarisation curve obtained in the dark at 30h and 54h for the Fe(+)|Air and Fe(-)|20% $CO_2$ cultures respectively. **(b)** Power curves calculated from the polarisation curves in (a). **(c)** Polarisation curve obtained under illumination at 234h 187h and 162h for the Fe(+)|Air, Fe(-)|Air and Fe(-)|20% $CO_2$ cultures respectively. **(d)** Power curve calculated from the polarisation curves in (c). Polarisation curves were measured with the following resistors (in $M\Omega$): 33, 10, 5.1, 2, 1, 0.56, 0.3, 0.1, 0.01. Error bars show $\pm1$ SEM of 3 independent replicates for each condition.

**Table S.3** – Parameters from the polarisation and power curves shown in Fig. S.10. Errors for $OCP$, $P_{max}$ and $J_{max}$. are ±1 SEM of 3 independent replicates. OCP was measured at t=0h (after four days of biofilm formation) before connection of the 33 $M\Omega$ external resistors. For each polarisation curve, $R_{int}$ was estimated from the slope of the linear portion of the curve fitted using Matlab's curve fitting toolbox, and values are quoted ±95% confidence interval of the gradient of the linear fit.

| **Condition** | $OCP$ $[mV]$ | $P_{max}$ $[\mu W \cdot m^{-2}]$ | $J_{max}$ $[\mu A \cdot m^{-2}]$ | $R_{int}$ $[M\Omega]$ |
|---|---|---|---|---|
| *Dark* | | | | |
| Fe (+) \| Air | ND | 0.32±0.15 | 15.0±5.63 | 5.27±8.7 |
| Fe (-) \| 20% $CO_2$ | ND | 21±1.4 | 569±51.6 | 1.47±0.081 |
| *Light* | | | | |
| Fe (+) \| Air | 44.9±7.4 | 9.8±2.7 | 458±112 | 0.76±0.34 |
| Fe (-) \| Air | 109±11.2 | 19±2.4 | 656±71.7 | 1.69±0.080 |
| Fe (-) \| 20% $CO_2$ | 127±8.8 | 20±1.4 | 467±29.1 | 1.34±0.036 |

**Abbreviations:** $OCP$ - Open Circuit Potential; $P_{max}$ - Maximum power; $J_{max}$ - Maximum current density; $R_{int}$ - Internal resistance; ND - Not determined.

### Derivatives of current density profiles

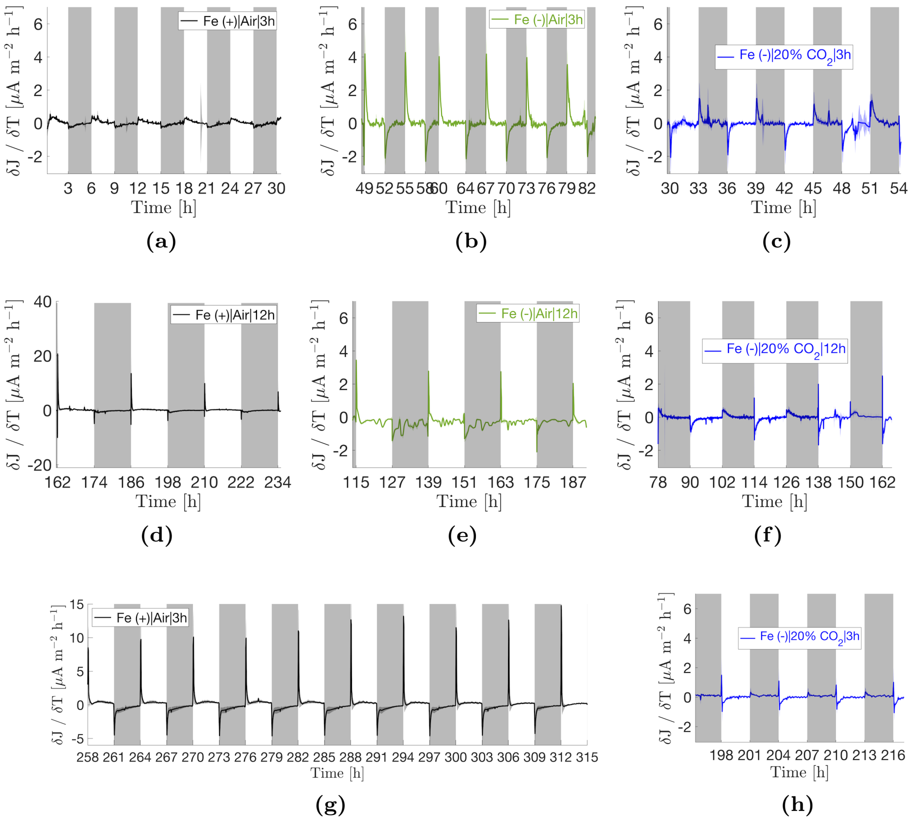

**Figure S.11 – Derivatives of current density profiles shown in Fig. 2 in the main text.** Each profile shows the mean of three independent replicates ± 1 standard error of the mean (shaded areas)

### Empirical Mode Decomposition of media only devices current density profiles

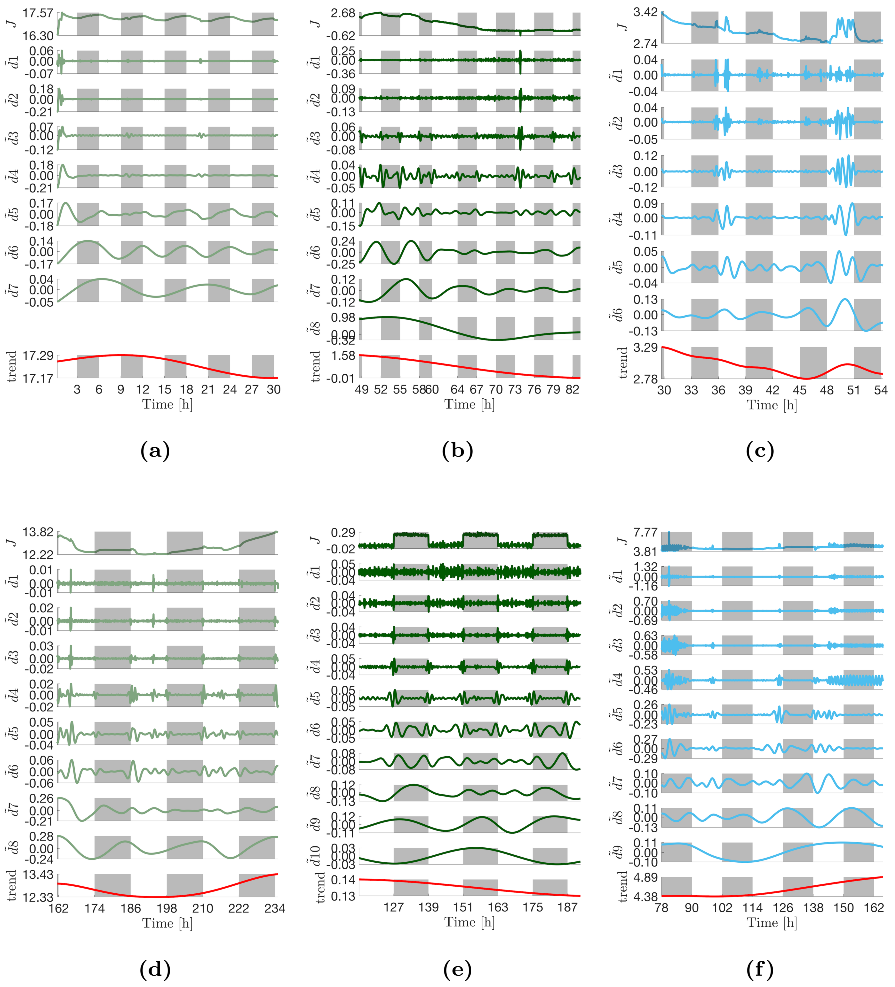

**Figure S.12 – Intrinsic mode functions (IMFs) extracted via the ICEEMDAN algorithm for media only devices**. **(a)** Fe (+)|Air|3h:Media. **(b)** Fe (-)|Air|3h:Media. **(c)** Fe (-)|20% $CO_2$|3h.:Media **(d)** Fe (+)|Air|12h:Media. **(e)** Blank device (no media, MEA only). **(f)** Fe (-)|20% CO2|12h:Media. For each decomposition, the top panel shows the mean current density profile, $\bar{J}$, as reported in Fig. 2 in the main text, the central panels show the extracted IMFs, and the bottom *trend* panel shows the final residue from the decomposition process $r8$ (red line). There was an unexpected positive dark response in the current density profiles, most clearly seen in (a) and (d). This was found to be due to an increase in electrical noise from the experimental rig's electrical connections when the light was turned off and a decrease when the light was turned on as measured in a blank device with an MEA only (no media) shown in (e).

### Hilbert spectra media only devices

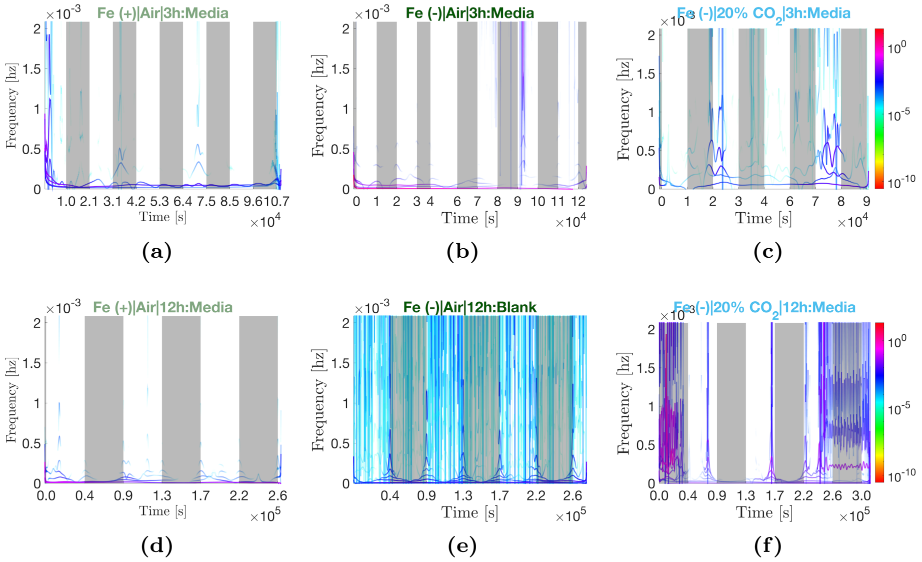

**Figure S.13 – Hilbert spectra for media only devices**. The spectra correspond to the IMFs shown in Fig. S.12.

### Marginal spectra media

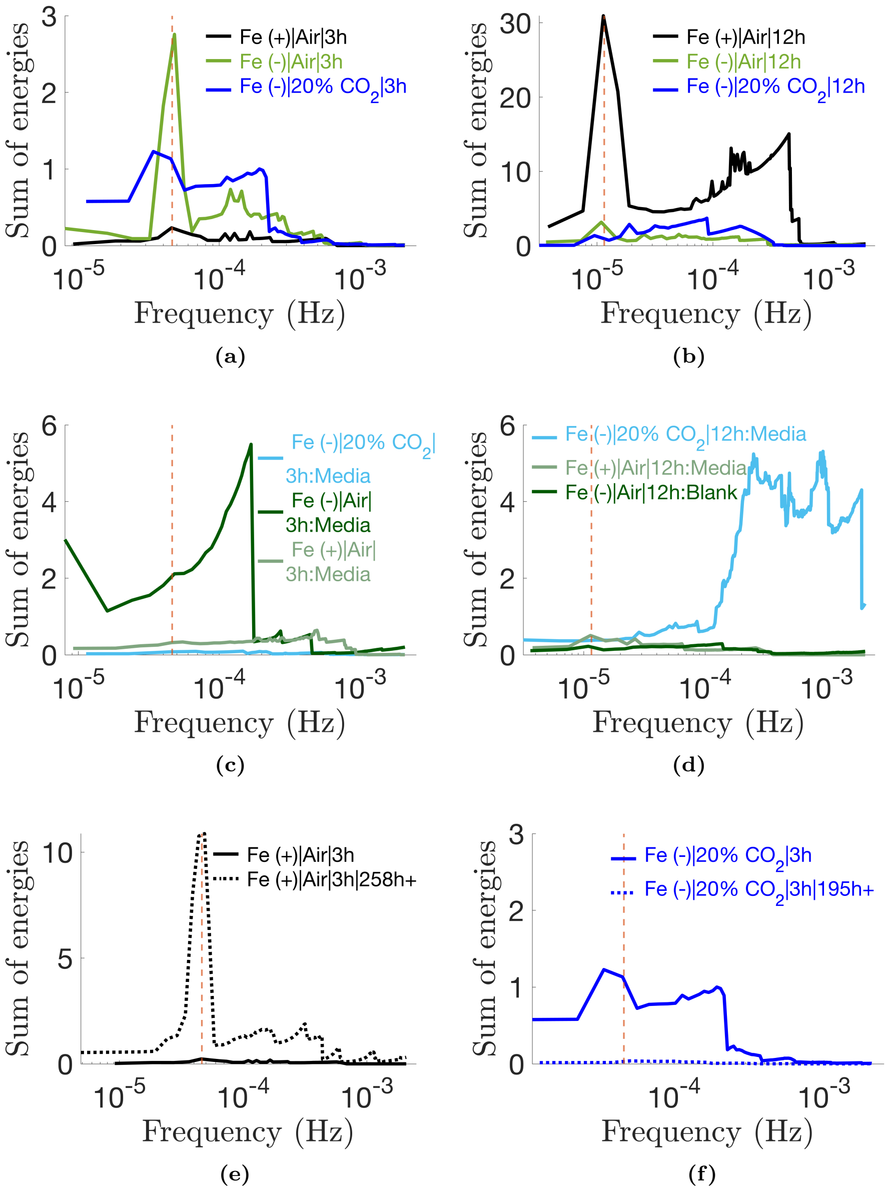

**Figure S.14 – Marginal spectra.** The marginal spectra show the spread of energies across the frequency range as calculated by Eq. 10 in the main text.
